# ERAP2 increases the abundance of a peptide submotif highly selective for the Birdshot Uveitis-associated HLA-A29

**DOI:** 10.1101/2020.08.14.250654

**Authors:** W.J. Venema, S. Hiddingh, J.H. de Boer, F.H.J. Claas, A Mulder, A.I. Den Hollander, E. Stratikos, S. Sarkizova, L.T. van der Veken, G.M.C. Janssen, P.A. van Veelen, J.J.W. Kuiper

## Abstract

Birdshot Uveitis (BU) is a blinding inflammatory eye condition that only affects HLA-A29-positive individuals. Genetic association studies linked *ERAP2* with BU, an aminopeptidase which trims peptides before their presentation by HLA class I at the cell surface, which suggests that ERAP2-dependent peptide presentation by HLA-A29 drives the pathogenesis of BU. However, it remains poorly understood whether the effects of ERAP2 on the HLA-A29 peptidome are distinct from its effect on other HLA allotypes. To address this, we focused on the effects of ERAP2 on the immunopeptidome in patient-derived antigen presenting cells. Using complementary HLA-A29-based and pan-class I immunopurifications, isotope-labelled naturally processed and presented HLA-bound peptides were sequenced by mass spectrometry. We show that the effects of ERAP2 on the N-terminus of ligands of HLA-A29 are shared across endogenous HLA allotypes, but discover and replicate that one peptide motif generated in the presence of ERAP2 is specifically bound by HLA-A29. This motif can be found in the amino acid sequence of putative autoantigens. We further show evidence for internal sequence specificity for ERAP2 imprinted in the immunopeptidome. These results reveal that ERAP2 can generate an HLA-A29-specific antigen repertoire, which supports that antigen presentation is a key disease pathway in BU.

## Introduction

Birdshot uveitis (BU) is a rare form of uveitis characterized by distinctive inflammatory foci across the retina, hypopigmented choroidal lesions, and cystoid macular edema, which causes visual impairment when undertreated^1,2^. Infiltration of T cells and elevated levels of T cell cytokines in eye tissues of patients suggest that T cell-mediated inflammation is among the driving disease mechanisms^3–6^. This is further supported by the fact that all patients with BU carry at least one copy of the Human leukocyte antigen (HLA)-A*29 allele, now widely considered as a prerequisite for diagnosis^7,8^. How HLA-A29 contributes to BU has remained unsolved, however, genetic association studies identified that in addition to the extreme association with the HLA-A*29:02 allele, polymorphisms in endoplasmic reticulum aminopeptidase (ERAP)-1 and ERAP2 confer strong disease risk^9,10^. Within the endoplasmic reticulum, ERAP aminopeptidases destroy or trim peptides to a length that is considered to influence their binding to HLA class I and presentation at the cell surface^11^. Importantly, of the two major haplotypes of ERAP2, the haplotype associated with canonical full-length ERAP2 (termed Haplotype A) is associated with BU^9^. The other common haplotype (haplotype B) encodes a transcript that undergoes alternative splicing and nonsense-mediated RNA decay, resulting in undetectable ERAP2 protein^12^. Because the risk haplotypes of ERAP genes for BU have been shown to result in lower cellular expression and activity of ERAP1 in combination with high cellular expression of functional ERAP2^10^, it is likely that ERAP2 generates a so far unknown, but highly HLA-A29-restricted antigen repertoire that dictates T cell- or NK cell responses. This renders antigen processing and presentation a key disease pathway in BU^9^.

ERAPs can trim the N-terminal residues of peptide substrates by sequestering the entire substrate inside the enzyme’s cavity where the sum of interactions of amino acid side chains are considered to determine the rate and outcome of peptide proteolysis^13,14^. Both ERAP1 and ERAP2 have been shown to have preferences for the internal sequence of the peptide, although these preferences are broad and no specific motif has been identified^13–16^. However, some studies have also shown that ERAPs can also trim peptide while they are bound onto MHC-I^17,18^, although this mechanism has been brought into question^19^. These and other observations^19^ support that ERAPs modulate a significant fraction of the ‘free’ peptide cargo before binding to HLA, which suggests that physiologically-relevant sequence specificities for ERAP2 may be deciphered from the presented peptide repertoire.

Mass-spectrometry based peptidomic studies of model high-passage cell lines have revealed that ERAPs can influence the peptide repertoire presented by HLA-A29^20,21^. However, to date, no studies have been conducted that studied the interaction of the major genetic risk haplotypes for ERAP1, ERAP2, and HLA-A*29:02 simultaneously in patient-derived tissues and compared the effects of ERAP2 on HLA-A29 to the other competing alleles expressed by the same cell. Knowing the potential effects of ERAP2 across HLA class I alleles is important to be able to separate potential disease effects from canonical antigen processing in studies of the immunopeptidome and may help predict the outcome of pharmacological interference of ERAP2 activity using small molecule inhibitors^22^.

We generated patient-derived lymphoblastoid cells that naturally express high levels of HLA and ERAPs, in which we stably expressed an autoantigen for BU (i.e. the retinal S-antigen, which is only expressed in the retina) to study if can present autoantigen fragments via HLA-A29. An advantage of using lymphoblastoid cells is that they express high levels of the immunoproteasome (e.g., LMP7 subunit)^23^, which is also highly expressed in photoreceptors of the retina where the immunoproteasome is essential for the maintenance of normal retinal function and vision transduction^24^. The use of newly established low-passage patient-derived antigen presenting cell lines better preserves the genetic architecture critically involved with BU in the context of physiologically relevant antigen processing.

In this study, we compared the immunopeptidomes of ERAP2-wild-type and ERAP2-deficient cells using mass spectrometry profiling of elutions from immunopurification with an HLA-A29-binding antibody and subsequent pan-class I antibody. Using several unbiased computational analyses, we accurately dissect the immunopeptidomes of HLA-A29 and other allotypes, which revealed commonly shared effects on position (P)1 and P7 of peptides across alleles, and hitherto unknown, specific effects on P2 in the HLA-A29 immunopeptidome with potential implications for the disease mechanisms of BU.

## Materials & Methods

### Generation of patient-derived EBV-immortalized B cell lines

EBV-immortalized lymphoblastoid cell lines (EBV-LCL) were generated from peripheral blood mononuclear cells (PBMC) from Birdshot patients, from which we selected a cell-line from a female patient (80 years old during sampling) homozygous for the risk haplotypes for *ERAP1* (*Hap10/Hap10*) and *ERAP2* (*HapA/HapA*)^10^. B95-8 marmoset-derived EBV supernatant was a kind gift from Dr. Willemijn Janssen, Center for Translational Immunology, UMC Utrecht. Cryopreserved PBMC were thawed and the cell number was determined. In a 24-well plate, 5-10^6^ cells were plated and cultured in freshly thawed EBV supernatant overnight at 37°C, 5% CO_2_. The next day, transformation-medium (RPMI 1640 + 10% FBS + 1μg/ml cyclosporine) was added into the wells. The EBV-infected cells were observed under the microscope to look for transformed LCLs in clusters. Patient-derived cell lines were cultured in Roswell Park Memorial Institute 1640 medium (RPMI 1640, Thermo Fisher Scientific) supplemented with 10% heat-inactivated fetal bovine serum (FBS, Biowest Riverside) and 1% penicillin/streptomycin (Thermo Fisher Scientific). To obtain stable cell lines overexpressing S-antigen, EBV-LCLs were transduced with the concentrated lentiviral supernatants (see **Supplemental Info**).

### ERAP2 KO using CRISPR-Cas9

For the generation of ERAP2 KO EBV-LCLs the Alt-R CRISPR-Cas9 system (Integrated DNA Technologies) was used and cells were electroporated with the Neon Transfection System (Thermo Fisher Scientific). First, the RNP complex was assembled by combining the crRNA CTAATGGGGAACGATTTCCT with the Alt-R tracrRNA (at a ratio of 1:1) and incubated at 95°C for 5 min, cooled down at room temperature and mixed with the Alt-R S.p Cas9 Nuclease and Buffer R (Neon system). After incubating the RNP complex for 10 minutes at room temperature, 8×10^5^ EBV-LCLs were mixed with the crRNA:tracrRNA-Cas9 complex and electroporated with two pulses of 1100 V and 30 ms each using the 10 μl Neon pipette tip. Electroporated cells were immediately taken up in antibiotic-free medium and cultured for minimal 7 days. This procedure was repeated for 3 times before ERAP2 protein expression levels were analyzed by western blot. A total of 5 rounds was required to reduce to levels of ERAP2 expression to near undetectable levels (**Figure 1B**).

**Figure 1.**
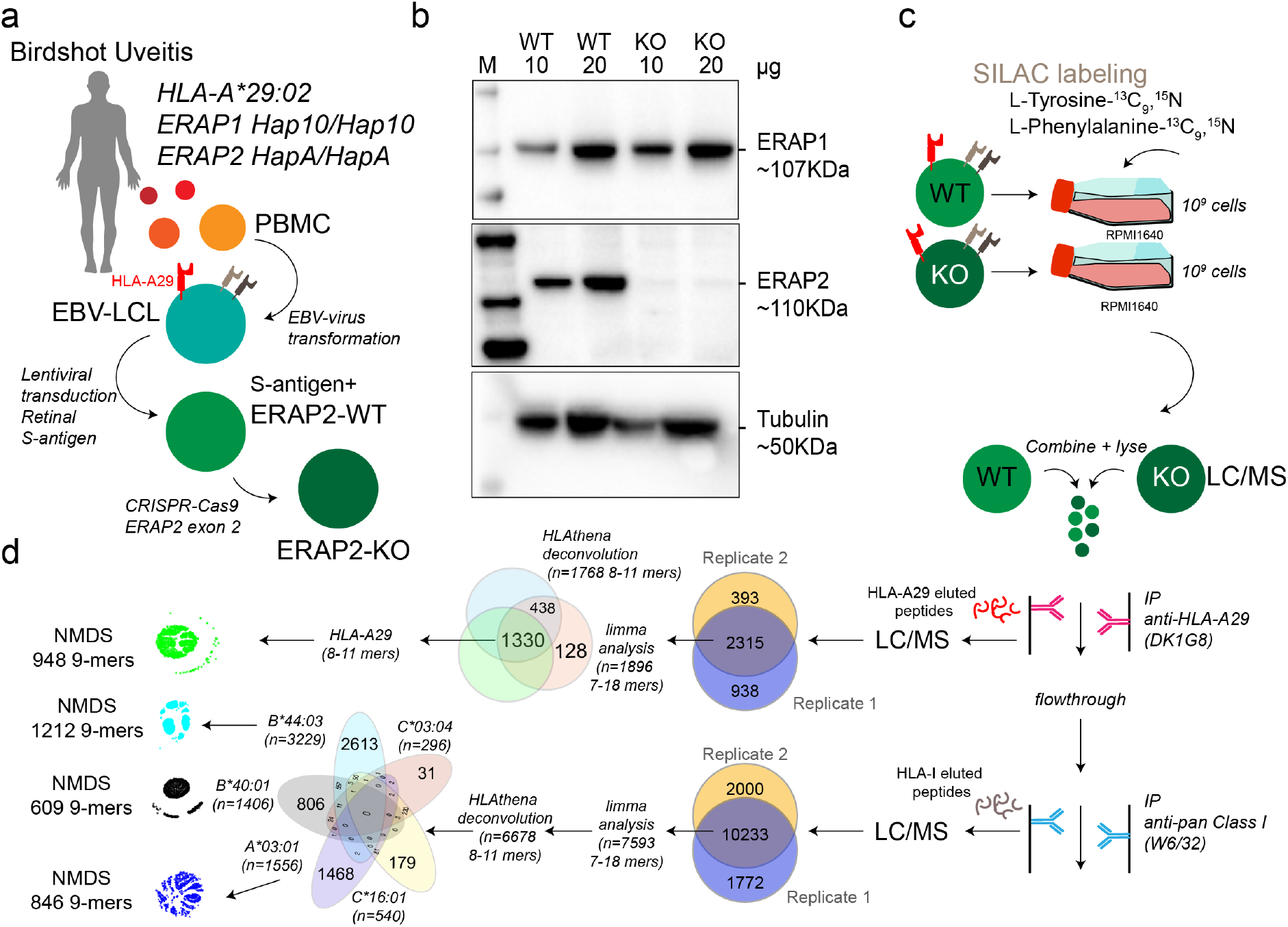
Study design and sample preparation. **a**) Design of the patient-derived model for antigen processing by ERAP2. **b**) Western blot analysis of the protein expression of ERAP1, ERAP2, and Tubulin as a control in the *HLA-A*29:02*-positive Birdshot uveitis model cell lines in ERAP2-wild type cells (WT) and cells after CRISPR-Cas9 mediated knock-out (KO) of ERAP2. The relative amount of protein (in microgram) used for each lane is indicated. M; marker. **c**) Overview of cultured SILAC labeled WT LCLs and unlabeled ERAP2 KO LCLs followed by combining the differentially labeled conditions for lysis and immunoprecipitation of HLA-A29 and, subsequently other HLA class I molecules, respectively. HLA-bound peptides were eluted, followed by LC/MS analysis. All steps in *c* were conducted in two separate experiments to generate biological replicates. **d**) Schematic overview of filtering steps of the identified peptides in this study. All peptides identified in both biological replicates with high confidence were filtered for *limma* analysis (see methods). After differential expression analysis, 8-11 mers were used to deconvolute and assign peptides to HLA alleles using *HLAthena*. The venn diagrams indicate the overlap from data sets and subsetting for subsequent analysis.

### Cell Culture and HLA-Peptide Immunopurification

For stable isotope labeling by amino acids in cell culture (SILAC), EBV-LCLs were cultured in customized RPMI with the same formula but lacking the two amino acids tyrosine and phenylalanine (Thermo Fisher Scientific) and with dialysed FBS (Thermo Fisher Scientific) in order to avoid unlabeled (i.e., ‘light’) amino acid carry-over. The medium was supplemented with L-Tyrosine-^13^C_9_,^15^N (Sigma Aldrich) and L-Phenylalanine-^13^C_9_,^15^N (Cortecnet). Wildtype EBV-LCLs were cultured with the customized medium (‘heavy’ labeled) and ERAP2-KO EBV-LCLs were cultured in RPMI with 10% non-dialyzed FBS (‘light’, without the labeled amino acids). Two independent experimental cultures were performed; Biological replicates were defined as two separate experiments starting from the CRISPR-Cas9-mediated ERAP2-KO (i.e., independent SILAC-cultures, immunopurification, elution and mass spectrometry profiling). In each experiment, cells from each condition were cultured to obtain 1×10^9^ cells in total per cell line. Cell pellets were stored at −20°C before mass spectrometry was performed. HLA class I molecules were isolated using standard immunoaffinity purification (IP) as described before^25^ from a fixed sample volume of 2.0 ml cell pellet per condition and biological replicate. IP was done using the human monoclonal antibody (mAb) DK1G8 (IgG1)^26^ derived from a HLA-A29-negative multiparous woman sensitized to HLA-A29 due to pregnancy, which specifically binds to 63-L-63-Q epitope in *HLA-A*29:01* and *A*29:02* and the very rare allele *A*43:01*, in a single antigen bead test. (https://www.epregistry.com.br/index/databases/database/ABC/), and a pan-HLA class I-specific mAb W6/32. Cell pellets from light and heavy labeled cell lines (ERAP2-WT and ERAP2-KO conditions) were combined and stored at −80 °C until mass spectrometry analysis.

### HLA-A29-binding and W6/32 antibodies

The hybridoma cell line producing HLA-A29-binding mAb DK1G8 was cultured in protein-free hybridoma medium supplemented with penicillin/streptomycin and L-glutamine in roller bottles. Cell culture supernatant was treated with Protein-A Sepharose beads to capture the mAb and eluted with glycine pH 2.5. Eluted mAb was covalently bound to Protein-A with dimethylpimelimidate for use in an immunoaffinity column (HLA-A29-Protein-A, W6/32-Protein-A Sepharose at 2.5 mg/ml). The columns were stored in PBS pH 8.0 and 0.02% NaN3 at 4 °C. HLA-bound peptides were extracted as described previously^25^.

### Isolation of HLA Class I presented Peptides

The extraction of peptides associated with HLA class I molecules was performed as described elsewhere^25^. Briefly, pellets from a total of 2 × 10^9^ LCLs were lysed for 2 hours at 4 °C in 50 mm Tris-HCl, 150 mm NaCl, 5 mm EDTA, and 0.5% Zwittergent 3-12 (N-dodecyl-N,N-dimethyl-3-ammonio-1-propanesulfonate) (pH 8.0) and the presence of Complete^®^ protease inhibitor (Roche). The preparation was centrifuged for 10 min at 2500 rpm and 4 °C and supernatant was transferred to a new tube and centrifuged for 40 min at 30,000 x g and 4 °C. The supernatant was pre-cleared with a 2-ml CL4B column and subjected to the immunoaffinity column (2ml with 5 mg ml). After washing, bound HLA class I-peptide complexes were eluted from the column and dissociated with 10% acetic acid. Peptides were separated from the HLA class I molecules via passage through a 10 kDa membrane (Microcon YM-10). The filtrate was freeze dried, dissolved in 50mM NH4HCO3 pH 8.4 and the peptides were further purified via ‘high pH reverse phase’ fractionation on a C18 column (Oasis HLB, Waters, Milford, MA). The peptides were eluted from the C18 Oasis column with successively 400 μl 10/90/0.1, 20/80/0.1 and 50/50/0.1 water/acetonitrile (ACN)/formic acid (FA), v/v/v.

### MS analysis

Peptides were lyophilized, dissolved in 95/3/0.1 v/v/v water/acetonitrile/formic acid and subsequently analysed by on-line C18 nanoHPLC MS/MS with a system consisting of an Easy nLC 1200 gradient HPLC system (Thermo, Bremen, Germany), and a LUMOS mass spectrometer (Thermo). Fractions were injected onto a homemade precolumn (100 μm × 15 mm; Reprosil-Pur C18-AQ 3 μm, Dr. Maisch, Ammerbuch, Germany) and eluted via a homemade analytical nano-HPLC column (30 cm × 50 μm; Reprosil-Pur C18-AQ 3 um). The gradient was run from 2% to 36% solvent B (20/80/0.1 water/acetonitrile/formic acid (FA) v/v) in 120 min. The nano-HPLC column was drawn to a tip of ~5 μm and acted as the electrospray needle of the MS source. The LUMOS mass spectrometer was operated in data-dependent MS/MS mode for a cycle time of 3 seconds, with a HCD collision energy at 32 V and recording of the MS2 spectrum in the orbitrap. In the master scan (MS1) the resolution was 60,000, the scan range 300-1400, at the standard AGC target @maximum fill time of 50 ms. Dynamic exclusion was after n=1 with an exclusion duration of 20s. Charge states 1-3 were included. For MS2 precursors were isolated with the quadrupole with an isolation width of 1.2 Da. Precursors of charge 1 were selected in the range of 800-1400, precursors of charge 2 were selected in the range 400-800, and precursors of charge 3 were selected in the range 300-600. The first mass was set to 110 Da. The MS2 scan resolution was 30,000 at the standard AGC target of 50,000 @dynamic injection time.

In a post-analysis process, raw data were first converted to peak lists using Proteome Discoverer version 2.1 (Thermo Electron), and then submitted to the Uniprot Homo sapiens canonical database (67911 entries), using Mascot v. 2.2.07 (www.matrixscience.com) for protein identification. Mascot searches were with 10 ppm and 0.02 Da deviation for precursor and fragment mass, respectively, and no enzyme was specified. Methionine oxidation was set as a variable modification.

### Differential expression analysis

Peptide confidence False Discovery Rates (FDRs) were calculated with the Mascot Percolator^27^ plug-in in Proteome Discoverer version 2.1 (Thermo Electron) and we used a strict target FDR of 1% (q<0.01) to obtain peptides detected with high confidence. To retrieve labeled peptides for downstream analysis, the high confidence peptides were further filtered to remove peptides with flags “InconsistentlyLabeled”, “NoQuanValues”, “Redundant”, “IndistinguishableChannels”. To detect significant changes in ligand abundance, we used the empirical Bayes workflow for mass spectrometry data based on the *limma28* and *qvalue29* R packages following Kammers and associates^30^ (see **Supplemental Info**). The *qvalue* R package was used to provide an unbiased estimate of the false discovery rate (FDR). Changes in peptide abundance between light and heavy conditions below a moderated q<0.01 (i.e., 1% empirical FDR) was considered affected by ERAP2. After differential expression analysis, peptides were assigned to HLA alleles using the *HLAthena* algorithm^31^, a state-of-the-art neural-network prediction algorithm trained on mass-spectrometry derived peptides from 95 mono-HLA expressing cell lines, which provides the binding score metric ‘MSi’ for each peptide and corresponding allele (range [0,1], MSi >0.6 was considered good, MSi>0.8 was considered strong). We used the GibbsCluster 2.0 server^32^ to deconvolute the detected 9-mers into a deconvolution solution of maximum three clusters (seeds=5, **λ**=0.7, σ=5, t=3). We picked a three-cluster solution that best matched the canonical binding motifs of the *HLA-A* alleles *HLA-A*29:02* (P**Ω**-Tyr/Y or Phe/F) and *HLA-A*03:01* (P**Ω**-Lys/K or Arg/R). For comparison of the effects of ERAP2 and ERAP1 on the HLA-A29 immunopeptidome, we used the 974 HLA-A29-presented peptides detected in both (identical peptide sequences) datasets from *Sanz-Bravo et al., 2018* (ERAP2, n=1140)^20^ and *Alvarez-Navarro et al., 2015* (ERAP1, n=5584)^21^ of which 917 showed normalized intensity values >0. In these studies, the normalized intensity ratio [IR] of each peptide in two cell lines was used to infer the relative abundance of each peptide in ERAP positive versus ERAP negative cell lines. The SWEIG cell line has very low ERAP1 levels and was considered functionally ‘negative’ for ERAP1. We plotted the normalized intensity ratio for each peptide as reported in the supplemental data from each study in **Figure 3I**.

### Non-metric mutidimensional scaling of peptides

Non-metric multidimensional scaling of 9-mers using entropy-weighted (*MolecularEntropy()* function from *HDMD* R package^33^ peptide distances in two-dimensional space was conducted following the method of Sarkizova and associates^31,34^. This method uses a Hamming distance calculated with an amino acid substitution matrix (adapted from Kim *et al.^35^*) that is inversely weighted according to positional entropy to obtain the pairwise “distance” between 9-mers. To map the peptide distances in two dimensions, for each analysed HLA allele, non-metric multidimensional scaling (NMDS) was used with 10 separate ordinations of 500 iterations using the *nmds()* function from the *ecodist* R package^36^. The configuration with the least stress was used for visualization of the peptidome. We next used *density-based spatial clustering of applications with noise* (DBSCAN)^37^ within the *fpc* R package^38^ to cluster peptides using the elbow method (*KNNdisplot function()* in *dbscan* R package^37^ to estimate the number of clusters that fit the data. Sequence logo plots were generated using the *ggseqlogo* R package^39^. The positional amino acid usage differences were calculated by determining the count for each amino acid at indicated positions (e.g., P1, P2) in the peptides using the *MolecularEntropy()* function from the HDMD *R* package and a fisher exact test was used (*fisher.test()* function in r base) to assess the differences at indicated positions. A chi-squared test (*chisq.test()* in r base) was used to assess for differences in the number of ERAP2 affected peptides per cluster. All *P* values were adjusted (termed *Padj*) using the bonferroni method as indicated. A *grand average of hydropathicity* (GRAVY) hydrophobicity index on the Kyte-Doolittle scale for each peptide was calculated with the *hydrophobicity()* function in *Peptides* R package^40^. Differences in binding scores and hydrophobicity index were assessed using the *dunnTest()* function in the *FSA* R package^41^.

### Western Blot analysis

Protein levels of S-antigen, ERAP1 and ERAP2 were analysed using western blotting. Total cell lysates were prepared using the NP40 lysis buffer (1% NP40, 135 mM NaCl, 5 mM EDTA, 20 mM Tris-HCl, pH=7.4) complemented with complete protease inhibitor cocktail (Roche). Protein lysates (10 μl/lane) were separated on a 4-20% Mini-PROTEAN TGX gel (Bio-Rad Laboratories) and transferred to a polyvinylidene difluoride membrane (Immobilon-P PVDF, Millipore). Membranes were blocked in 5% nonfat dry milk in TBST and probed overnight at 4°C with antibodies recognizing ERAP1 (AF2334, R&D Systems), ERAP2 (AF3830, R&D Systems), S-antigen (α-mGFP, TA180076, Origene, to detect the fusion protein S-antigen-GFP) or α-tubulin (T6199, Sigma). After washing, membranes were incubated with anti-mouse secondary antibody conjugated to horseradish peroxidase (HRP) (DAKO) or anti-goat secondary antibody conjugated to HRP (DAKO). Protein bands were detected with Amersham Prima Western Blotting (RPN22361, GE Healthcare) on the ChemiDoc Gel Imaging System (Bio-Rad Laboratories). The ratio of the intensity was calculated using Image Lab 5.1 (Bio-Rad Laboratories) for each experiment.

### High-density SNP-array analysis

SNP-array copy number profiling and analysis of regions of homozygosity were performed on DNA isolated from WT and CRISPR-Cas9 edited LCLs (ERAP2-KO) according to standard procedures using the Infinium Human CytoSNP-850K v1.2 BeadChip (Illumina, San Diego, CA, USA). Samples were scanned using the iScan system (Illumina). Subsequently, visualizations of SNP-array results and data analysis were carried out using NxClinical software v5.1 (BioDiscovery, Los Angeles, CA, USA). Human genome build Feb. 2009 GRCh37/hg19 was used.

### Data availability

Analysis code, genotype data, and supporting data files can be found at https://github.com/jonaskuiper/ERAP2_HLA-A29_peptidome. Mass spectrometric raw data has been deposited in the MassIVE depository (MassIVE dataset XXXXX) under the creative commons zero license (CC0 1.0).

## Results

### Generation of a model for ERAP2-mediated antigen processing and presentation

We generated lymphoblastoid cells (LCLs) from a *HLA-A*29:02-*positive Birdshot patient homozygous for risk haplotypes of *ERAP1* (*Hap10/Hap10*) and *ERAP2* (*HapA/HapA*)(**Figure 1A**)^9,10^ and the retinal S-antigen was stably expressed by lentiviral transduction (see **Supplemental Notes**). Genotyping of the patient revealed *HLA-A *29:02, HLA-A *03:01, HLA-B*40:01, HLA-B*44:03, HLA-C*16:01*, and *HLA-C*03:04* alleles. Because the risk allotype of ERAP1 shows relatively low aminopeptidase activity^10^, we focused our analysis on the effects of ERAP2 on the immunopeptidome. We used CRISPR-Cas9 ribonucleoprotein delivery with a guideRNA targeting exon 2 in *ERAP2* (**Figure 1A**) to disrupt protein expression of ERAP2 and generate an ERAP2-KO LCL, while preserving the protein expression of ERAP1 (**Figure 1B**). SNP-array copy number profiling and analysis of regions of homozygosity were performed using the Infinium Human CytoSNP-850K capable of detecting genomic gains and losses with an approximate resolution of ~10 kb by profiling 850,000 single nucleotide polymorphism (SNP) markers spanning the entire genome. SNP-array analysis resulted in a normal female array profile (arr(1-22,X)x2) for both cell conditions (**Figure S1**), and detected no changes between the WT and ERAP2-KO clones, including the ERAP region *5q15* (**Figure S2**). This confirms that our editing strategy did not introduce wide spread genomic changes and thus that the conditions are highly suitable for comparison. Genotype date for 92 SNPs at *5q15* for these cell lines is shown in **Table S1**.

Next, we used stable isotope labeling by amino acids in cell culture (SILAC) to incorporate “heavy” L-Tyrosine-^13^C_9_,^15^N (Tyr/Y) and L-Phenylalanine-^13^C_9_,^15^N (Phe/F) in the ‘wild type’ (WT) LCLs and compare these to unlabeled (“light”) culture conditions for the ERAP2-KO LCL cells (**Figure 1C, 1D**). The amino acids Y/F are observed in 95% of previously identified HLA-A29 ligands (**Figure 2A**), but are also found in the majority of peptides presented by the other HLA allotypes – with exception of HLA-B40:01.

**Figure 2.**
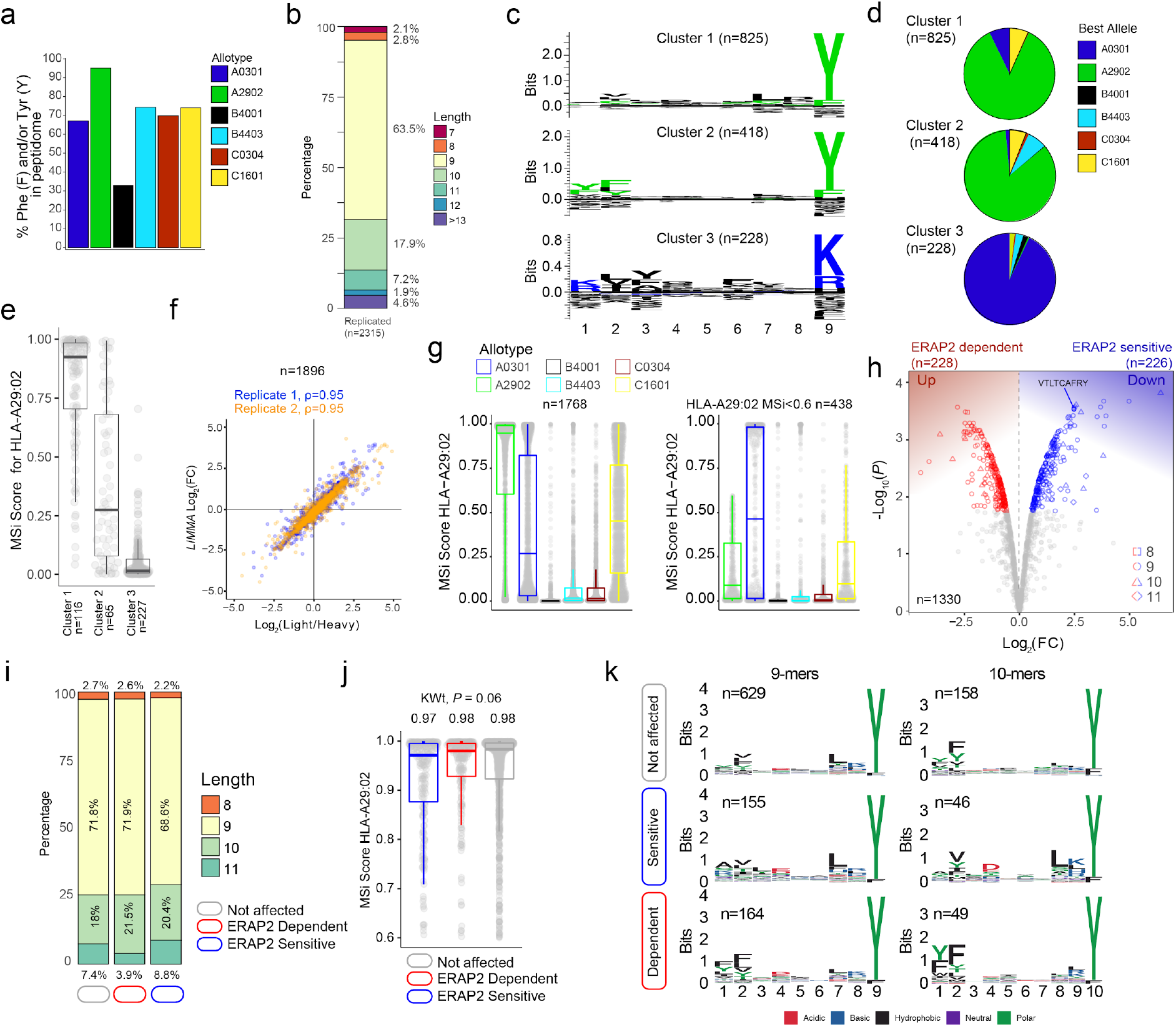
ERAP2 shapes the HLA-A29 peptidome. **a**) The percentage of peptides that contain Phenylalanine and/or Tyrosine in peptidomic studies of monoallelic cell lines by Sarkizova and associates^31^. **b**) The length distribution of the 2315 peptides detected in both biological replicates. c) *GibbsCluster 2.0* results for unbiased clustering of the 9-mers (n=1471 unique peptides) eluted with the HLA-A29-binding monoclonal antibody. The motifs correspond with the *HLA-A* genotype (*HLA-A*29:02/HLA-A*03:01*) of the sample. Cluster 1 and 2 match the binding motif of HLA-A29:02, and Cluster 3 matches the binding motif of HLA-A03:01. **d**) Pie diagrams (percentages) of best assigned alleles for the peptides in the clusters identified in *c*. The alleles which correspond to the best score for each peptide (‘Best Allele’ output from *HLAthena*) was used to obtain the percentages of peptides assigned to each of the six *HLA-A, -B*, and *-C* alleles. **e**) The binding scores for HLA-A29:02 for peptides from the clusters identified in *C* assigned to the other alleles. **f**) Strong correlation between the raw peptide abundance data (n=1896) and normalized data by *limma* used in the differential expression analysis. **g**) The 1768 8-11 mers before (left, plot) and after (right plot) filtering out the 1330 HLA-A29-binding peptides. **h**) Volcano plot of the differentially expressed 8-11 mers. In red are peptides that are increased in expression in the presence of ERAP2, while peptides indicated in blue are decreased. The identified peptide VTLTCAFRY from the retinal S-antigen is indicated. **i**) The length distribution and **j**) binding scores for HLA-A29 of the peptide groups identified in *h*. **k**) Sequence logos generated using a non-redundant list of 9-mers and 10-mers (11-mers see **Figure S4**).

### Capture of a high-quality HLA-A29 peptidome

Using the HLA-A29-binding antibody, a total of 2315 unique peptides were identified with high confidence (Mascot Percolator q<0.01) between biological replicates (*Jaccard* similarity = 0.64)(**Figure 1D**) that were used for further analysis. These were predominantly 9-11 mers (88%), which fits the length distribution^42^ of HLA-A29 ligands (**Figure 2B**). The HLA-A29-binding antibody may weakly cross-react with other HLA-A allotypes (see ***Methods***). This is of relevance given that *HLA-A *29* alleles are low expressed *HLA-A* alleles^43^ compared to high expressed *HLA-A*03* alleles. We used *GibbsCluster 2.0* for unbiased clustering of the peptides, which found a deconvoluted solution that consisted of three clusters; two motifs fitting the canonical HLA-A29:02 binding motif (C-terminal position Y or P**Ω**-Y) and one cluster highly similar to the dominant HLA-A03:01 motif (P**Ω** Lysine (K) or Arginine (R)(**Figure 2C**) and shows that the HLA-A29 antibody cross-reacts with HLA-A03:01. Indeed, when we used the *HLAthena* algorithm^31^, ligands in cluster 1 and 2 were predominantly assigned to HLA-A29:02 (84% and 86%, respectively), and 93% of ligands in cluster 3 were assigned to HLA-A03:01 (**Figure 2D**). However, because 66% and 20% of peptides in clusters 1 and 2 assigned to other endogenous HLA alleles also showed high binding scores for HLA-A29:02 (**Figure 2E**), we later choose to filter the dataset using bindings scores for HLA-A29:02 (**Figure 2G**).

Because we were interested in determining significant changes in peptide abundance associated with ERAP2, we first jointly analyzed the relative abundance (fold change) of light (KO) over heavy (WT) labeled peptides from both experiments using *limma*^30^. A total of 1,896 peptides (**Figure 2F**) were detected in both light and heavy channels and used for analysis. Analysis of peptides unique to one of the conditions is shown in the Supplemental Info. Note that the log fold changes of pooled normalized peptides abundances from light and heavy channels by *limma* strongly correlate (spearman r = 0.95) with the light/heavy ratio abundance of each experiment (**Figure 2F**), thus the normalization steps preserve the data structure, while improving the power to detect significant changes^30^. From the 1,330 8- to 11-mers HLA-A29 epitopes (MSi>0.6 by *HLAthena*)(**Figure 2G**), 1,195/1,330 (89%) of the peptides in our HLA-A29 dataset have been reported as ligands for HLA-A29 of which 78% detected in mono-allelic or homozygous HLA-A29-expressing cell systems^20,31^, supporting the notion that the approach taken yields an accurate representation of the peptide-presenting properties of HLA-A29:02.

### ERAP2 shapes P1 of HLA-A29 ligands

At a false discovery rate of 1%, in ERAP2-WT compared to ERAP2-KO cells, a total of 226 peptides were detected at decreased abundance in the binding groove of HLA-A29 (termed ERAP2-”sensitive” peptides), and 228 peptides were increased in abundance (termed ERAP2-“dependent” peptides) (**Figure 2H and Table S2**). We detected the 9-mer VTLTCAFRY from retinal S-antigen, which was ~6-fold higher (Log_2_[FC] = 2.45) in ERAP2-KO cells compared to ERAP2-WT cells, indicating ERAP2 destroys this epitope (**Figure 2H and Table S2**). We observed moderate changes in the length distribution (**Figure 2I and Figure S3**) or predicted binding affinities of peptides affected by ERAP2 (Kruskal-Wallis *P* = 0.06)(**Figure 2J**). In contrast, comparison of the peptide motifs revealed evident and consistent changes at the N-terminal amino acid positions for ERAP2-sensitive 9-11 mer peptides compared to peptides not affected by ERAP2 (**Figure 2K**, **Figure S4**), which aligns with the current view that ERAP2 trims the N-terminal amino acids of peptide substrates^13^. In detail, P1 of 9-mers revealed a contrasting residue preference for ERAP2-sensitive and ERAP2-dependent peptides (**Figure 3A**); Alanine(A), K, and R amino acids were seen significantly more often, while amino acids Y and F were seen significantly less often in sensitive peptides compared to nonaffected peptides (Fisher’s Exact test, *Padj* <0.05, **Table S3**). In contrast, the most common P1 residues for dependent and non-affected peptides were Y and F (Y/F at P1; 45% and 30%, respectively) with F statistically more abundant at P1 and P2 in dependent peptides (**Figure 3A**, **Table S3-4**). Intriguingly, we detected no significant effects of ERAP2 at the N-terminal residue of the precursor peptide (position P-1)(**Figure S5**). Together these data show that ERAP2 has a selective effect on P1 of the HLA-A29 immunopeptidome in part by driving the depletion of peptides with preferred P1 substrates (e.g., A, K, R)^44^ of ERAP2. This finding is consistent with previous reports that ERAP2 has primarily a destructive role by over-trimming susceptible peptide sequences and thus removing them from the immunopeptidome^44^.

**Figure 3.**
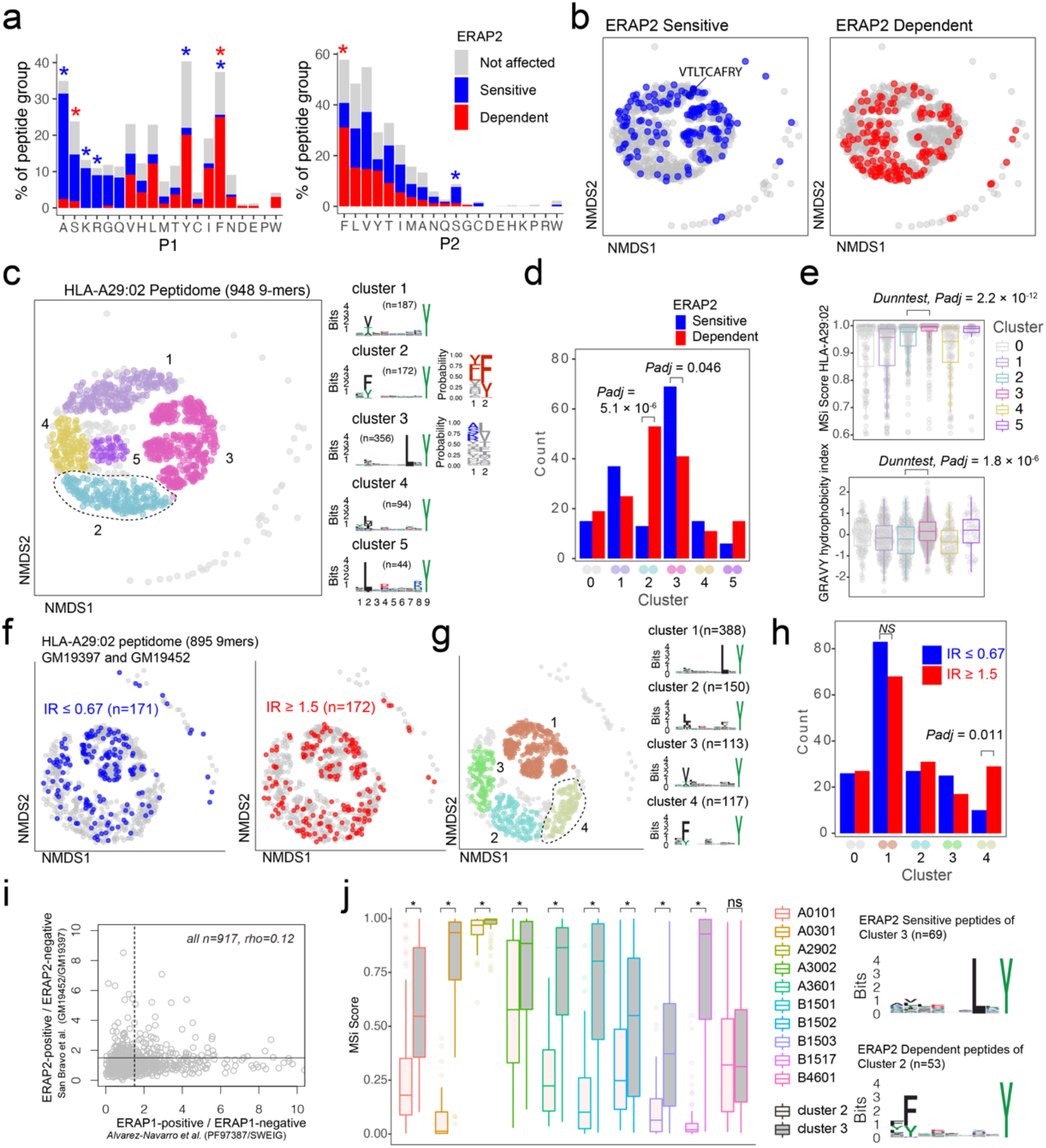
ERAP2 increases the abundance of a peptide-motif that is highly selective for HLA-A29. **a**) Comparison of amino acid proportion at P1 and P2 of 9-mers (% for each group) between peptides that decrease in abundance (‘sensitive’ peptides, significant changes indicated by the blue asterix), peptides that increase in abundance (‘dependent’ peptides, significant changes indicated with the red asterix), compared to peptides not affected in ERAP2-WT cells (in grey). Statistics from the fisher tests are indicated in **Table S3-4**. **b**) Non-metric multidimensional scaling (NMDS) of 948 9-mers from HLA-A29:02. Peptide distance was defined on the basis of sequence similarity. Each circle represents a unique peptide and is color-coded according to the effect of ERAP2; grey: not affected, blue: Peptides decreased (‘sensitive’) in abundance and in red peptides that increased (‘dependent’) in abundance in the ERAP2-WT condition compared to the ERAP2-KO condition. The peptide VTLTCAFRY from the retinal S-antigen is indicated. **c**) NMDS plot of HLA-A29:02 with 9-mer peptides color-coded according to the clustering by DBSCAN. Sequence logos and two probability plots representing these clusters are shown. **d**) Comparison of the number of ERAP2-sensitive and -dependent peptides in cluster. *Padj* = bonferroni corrected (5 clusters) *P* values from X^2^ tests. **e**) binding scores (MSi metric from *HLAthena*) for HLA-A29:02 and hydrophobicity index for each cluster. **f**) NMDS of 895 shared 9mers from HLA-A29-positive cell lines GM19452 (ERAP2-positive) and GM19397 (ERAP2-deficient) from *Sanz-Bravo* and associates^20^. In this study, the normalized intensity ratio (GM19452/GM19397) of each peptide in the two cell lines was used to infer the relative abundance of each peptide, which we adapted to assign peptides as ERAP2-sensitive (IR ≤ 0.67, n=171 peptides) or ERAP2-dependent (IR ≥ 1.5, 172 peptides). **g**) Four clusters were estimated by DBSCAN. **c**) **h**) Comparison of the number of ERAP2-sensitive and ERAP2-dependent peptides in each peptide cluster identified in *g* similar to *d*. **i**) Correlation plot of the effects of ERAP1 and ERAP2 on the HLA-A29 peptidome (see methods). The spearman’s correlation (rho) is shown for 947 HLA-A29-eluted peptides detected in two studies. The black lines indicate the threshold of the normalized intensity ratio >1.5 used in each of the studies. This analysis suggests very low correlation between the effects of ERAP1 and ERAP2 on similar peptides presented by HLA-A29. **j**) Predicted binding scores for ERAP2-dependent peptides in cluster 2, and ERAP2-sensitive peptides in cluster 3 for HLA-A29:02 and 9 HLA alleles with relatively similar binding motifs (based on Sarkizova *et al*.^31^). *) indicates bonferroni corrected *P*<0.05 from a Dunn’s Test.

### ERAP2 increases the abundance of peptides with a cryptic aromatic P2 motif

ERAP2 trims peptides by sequestering them into the relatively large internal enzyme cavity^13^, where peptide side chains across the amino acid sequence can interact with pockets inside the cavity of ERAP2^13,14^. To evaluate if sequence-specific selectivity^16^ by *ERAP2* could be interpreted from the HLA-A29 peptidome, we conducted non-metric multidimensional scaling (NMDS) of all 9-mers^31^. This analysis projects peptides in two-dimensional space based on the similarity of the amino acid sequences (**Figure 3B**). Considering peptides with significant changes between ERAP2-WT and -KO conditions revealed distinct patterns for co-clustered (“similar”) peptides, with ERAP2-sensitive peptides located ‘away’ from ERAP2-dependent peptides (**Figure 3C**). To quantify these differences, we compared the amount of ERAP2-sensitive (**Figure 3B** in blue, n=155) versus ERAP2-dependent peptides (in red, n=164) across five clusters of peptides or ‘submotifs’^34^. This analysis revealed that ERAP2-sensitive peptides were overrepresented in cluster 3 (X^2^, Bonferroni n=clusters, *Padj* = 0.046) and ERAP2-dependent peptides overrepresented in cluster 2 (*Padj* = 5.1 × 10^-6^)(**Figure 3D**). Cluster 2 (n=172 in total) was defined by nonpolar aromatic residues F (*Padj* = 1.0 × 10^-49^), or Y (*Padj* = 2.1 × 10^-22^) at P2 (F/Y in 97% of peptides in cluster 2 compared to 13% of peptides in all other clusters). ERAP2-dependent peptides (n=53) made up a considerable proportion of cluster 2 (unaffected peptides; n=106).

Peptides in cluster 3 (n=356) were distinguished by a L at P7 (99% of peptides in cluster 3 compared to 3% in other clusters, *Padj* = 2.0 × 10^-230^)(**Figure 3C and Table S5**). Peptides in cluster 3 showed an overall higher binding score for HLA-A29:02 and higher hydrophobicity index compared to cluster 2 (**Figure 3E**). Note that submotifs cluster 2 and cluster 3 are *bona fide* submotifs of HLA-A29 that are highly reproducible in other datasets (cluster 1 and 4 in **Figure 3F and G** and cluster 1 and 3 in **Figure S6**). We further replicated our findings in HLA-A29 immunopeptidome data from Sanz-Bravo *et al.^20^* of ERAP2-competent and naturally ERAP2-deficient HLA-A29-positive cell lines (**Figure 3H**) and, thus, demonstrate that ERAP2-positive cell lines commonly display selectively increased peptides with the motif of cluster 2. In contrast, ERAP1 did not selectively contribute to cluster 2 peptides (**Figure S6**). Also, the effect of ERAP1 and ERAP2 on HLA-A29 peptides correlated weakly (spearman rho=0.12, **Figure 3I**), suggesting non-redundant effects for ERAP1 and 2 on the HLA-A29 peptidome. Although the analysis for 10-mers (n=235) in our dataset was considered to lack sufficient resolution to map the effects of ERAP2 on the submotif level, most of the ERAP2-dependent 10-mers also mapped to a submotif of HLA-A29 with F at P2 (**Figure S7**). In summary, ERAP2 selectively increases the expression of HLA-A29-binding peptides with a submotif with aromatic residues at P2.

### ERAP2-dependent peptides of cluster 2 are selective for HLA-A29

HLA class I peptides display promiscuity^45^ and it is therefore of interest that HLA-A03:01 can present peptides with a Y at P**Ω** (similar to HLA-A29) only with L at P7 is present^42^. As expected, peptides from cluster 3 (**Figure 3C**) were also predicted as potential binders for HLA-A03:01, while cluster 2 peptides (**Figure 3C**) were not (**Figure S8**). To further test the HLA allotype restriction, we compared the binding scores for the differentially expressed peptides in cluster 2 and 3 for eight alleles which display binding motifs that overlap with HLA-A29:02 (based on Sarkizova *et al.^31^*). As shown in **Figure 3J**, ERAP2-sensitive peptides in cluster 3 show relatively good (MSi>0.8) binding scores for several other alleles (e.g., HLA-A30:02). Note that the S-antigen peptide VTLTCAFRY (in cluster 1, **Figure 3B**) also shows good binding scores for other alleles (e.g., HLA-A30:02 MSi = 0.86). In contrast, ERAP2-dependent peptides from cluster 2 are predicted to poorly bind the other class I alleles with an overall similar binding motif (median MSi<0.6)(**Figure 3J**), indicating that this cluster is highly specific for HLA-A29:02. We extended this analysis to 95 alleles, which supported that the ERAP2-dependent peptides in cluster 2 are highly specific for HLA-A29 (**Figure S9**), with the exception of *HLA-C*14:03* (>100 times lower allele frequency compared to *HLA-A*29:02* in European populations) (**Table S6**). The motif of cluster 2 peptides is present in the amino acid sequences of proteins encoded by ~300 genes highly expressed in the retina (**Table S7**), of which putative HLA-A29-restricted peptides (MSi>0.9 for HLA-A29 and MSi<0.6 for 94 other alleles) were found in key factors in melanocyte biology (ARMC9, OCA2, SLC45A2, PLXNC1)(**Table S8**). This is of significance, because progressive loss of ocular melanocytes is a hallmark feature of BU^2,5,8,53,54^. We conclude that these data support that ERAP2 may apply selective pressure on the repertoire of HLA-A29.

### ERAP2 has similar effects on P1 across the HLA class I immunopeptidome

Next, we were interested to see how ERAP2 affects the global peptidome of the other class I alleles. We use the flow-through of the HLA-A29-binding antibody immunopurifications to capture HLA class I molecules (**Figure 1D**). After filtering, a total of 10,233 unique peptides were identified between biological replicates (*Jaccard* similarity = 0.73) of which 6,678 8-11 mers were considered for differential expression analysis (**Figure 1D**)(**Table S9**). A total of 2,170 peptides were differentially expressed (**Table S9**). Notwithstanding allele-specific differences, K, R, and A were seen more often at P1 of ERAP2-sensitive peptides, while F and Y were typically underrepresented across the other five alleles (**Figure 4**)(**Tables S10-S14**). This was supported by a global assessment of all 9-11-mers (**Figure S10**). These results indicate that ERAP2 has globally similar effects on P1 across HLA allotypes and in line with the observation that the P1 across HLA class I ligands is enrichment for residues A, K, and R ^34^.

**Figure 4.**
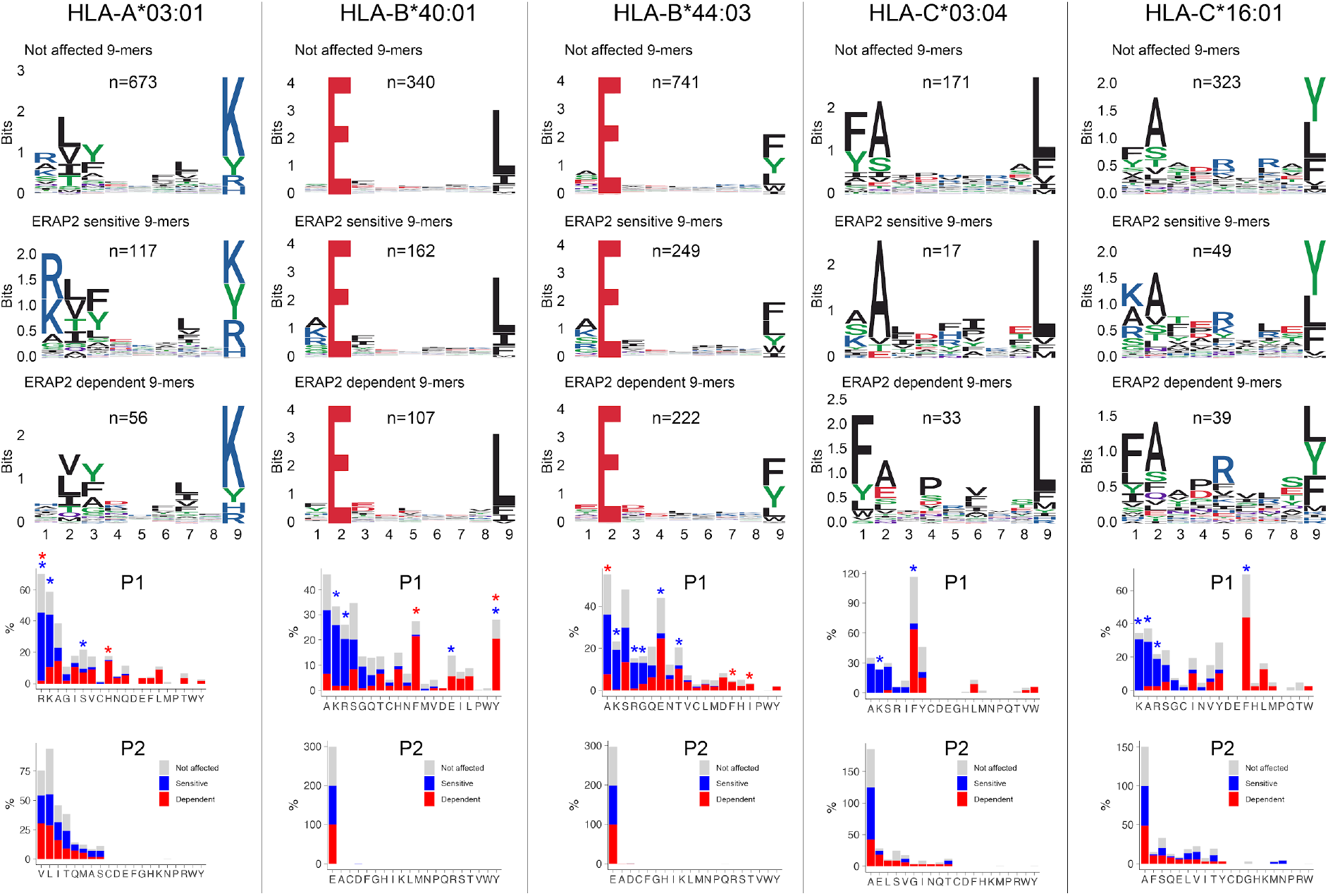
ERAP2 shapes P1 across the HLA class I immunopeptidome. Sequence motifs depict specific amino acid preferences at P1-P9 and were generated from a non-redundant list of 9-mers for each class I allele. Comparison of amino acid proportion at P1 and P2 of 9-mers (in percentage for each group of peptides) between peptides that decrease in abundance (‘sensitive’ peptides, significant changes indicated with the blue asterix), peptides that increase in abundance (‘dependent’ peptides, significant changes indicated with the red asterix), compared to peptides not affected in ERAP2-WT cells (in grey). The *P* values and summary statistics from the fisher tests are indicated in **Table S10-S14**.

### Internal sequence preferences of ERAP2 can be interpreted from the immunopeptidome

We further conducted NMDS of the 9-mers for HLA-A03:01, HLA-B40:01, and HLA-B44:03 (**Figure 5A, D, G** and **Figure S11**). The *HLA-C* peptidomes captured were too sparse to provide sufficient resolution (**Figure S12**). Investigation of HLA-A03:01 was hampered by a relatively high level of submotifs, characteristic for this allele^31,34^, in comparison to the density of the peptide data (**Figure 5B**), possibly due to loss of peptides by the initial immunopurification (**Figure 1D**). Regardless, ERAP2-sensitive peptides were enriched in cluster 4 (X^2^, *Padj* = 9.5 × 10^-3^)(**Figure 5C**), but 37/92 (40%) peptides of cluster 4 were also in the HLA-A29 peptidome (20 in cluster 3 of HLA-A29:02, **Figure 3C**). Reanalysis of immunopeptidome data from mono-allelic cell lines^31^ support that HLA-A29:02 and HLA-A03:01 can each present peptides with the motif of cluster 4 (**Figure S13**) and demonstrates that ERAP2 influences multiple alleles in part by peptide promiscuity. Considering the other clusters, no evidence for effects of ERAP2 beyond P1 could be observed.

**Figure 5.**
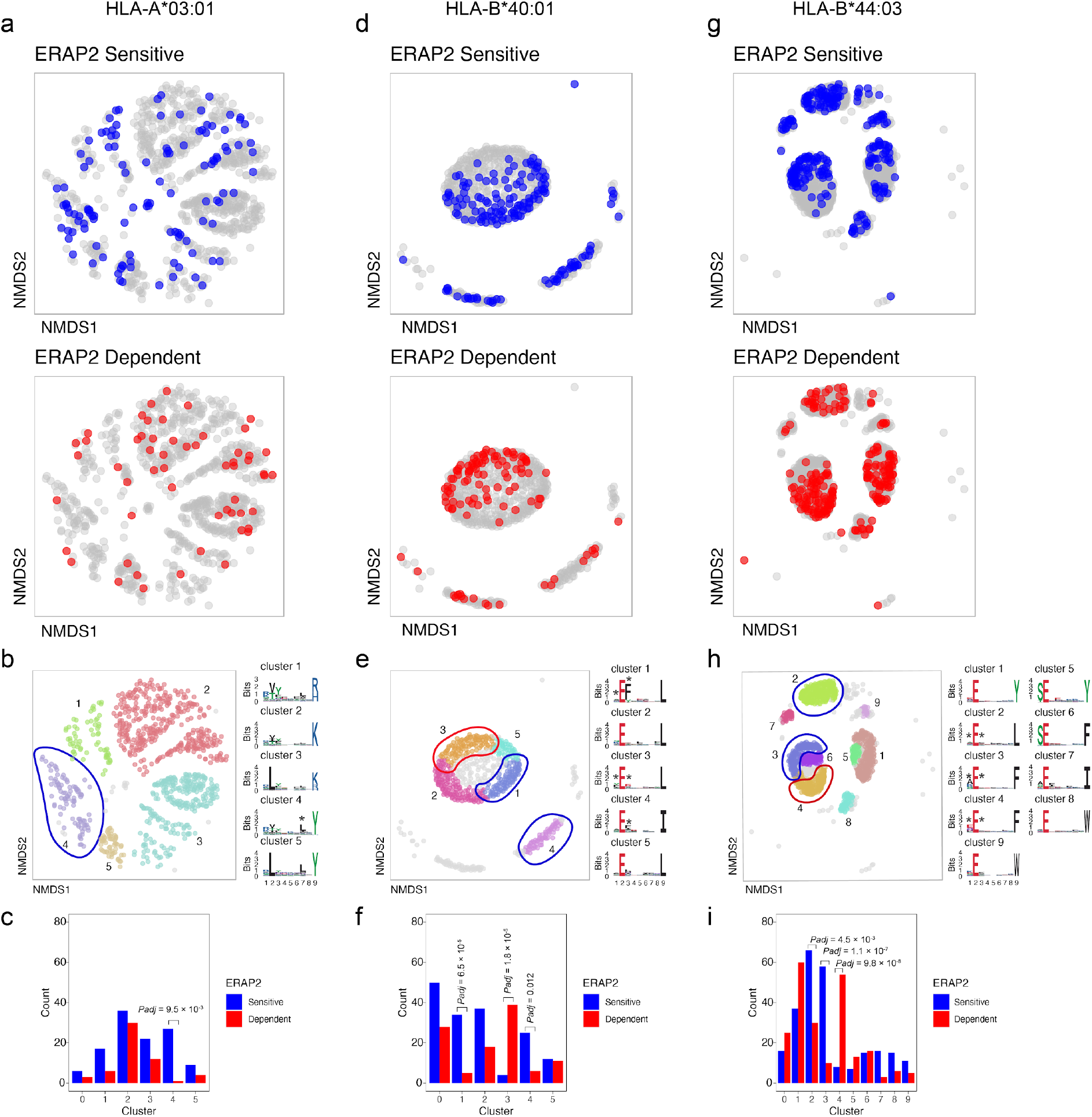
NMDS plots showing 9-mer peptide clustering for individual HLA alleles. Non-metric multidimensional scaling (NMDS) visualization of 9-mer peptides and ERAP2 affected peptides for *HLA-A*03:01* (**a,b,c**), *HLA-B*40:01* (**d,e,f**), and *HLA-B*44:03* (**g,h,i**). Peptide distance was defined on the basis of sequence similarity. Each circle represents a unique 9-mer peptide and is color-coded according to the effect of ERAP2; grey: not affected, blue: ERAP2-sensitive peptides red: ERAP2-dependent peptides. The NMDS plot of clusters of peptides for each class I allele peptide are color-coded according to the clustering by DBSCAN. Sequence logos representing these clusters are indicated. * indicates significant changes of amino acid composition tested at P1, P2 and and/or P7 (Fisher’s exact test corrected for 20 amino acid residues. Given the entropy-weighted clustering, anchor positions P2 and P9 were not considered for testing. Clusters with significant differences in the count of ERAP2-sensitive and -dependent peptides are highlighted with blue and red ellipses and correspond with the barplots in c,f, and i. The predicted binding scores for each cluster is shown in **Figure S11**. *Padj* = bonferroni corrected (n=clusters) *P* values from X^2^ tests. All other comparisons were *Padj>0.05*.

In contrast to HLA-A03:01, strong residue preferences at P2 and P**Ω** of HLA-B40:01 resulted in few submotifs (**Figure 5D**). The distribution of ERAP2-sensitive ‘away’ from dependent peptides in two-dimensional space was reminiscent of the ‘pattern’ observed in the projection of HLA-A29:02 peptides (**Figure 5D**). Submotif analysis revealed that cluster 1 and 4 were enriched for sensitive peptides and were distinguished by a preference for F or Y at P3 (**Figure 5E, F**)(**Table S15**). Cluster 3 (enriched for dependent peptides) was distinguished by a F/Y at P1 (**Figure 5E**), similar to the overall motif of ERAP2-dependent peptides.

Finally, HLA-B44:03 submotifs enriched for sensitive peptides (cluster 3 and cluster 2) (**Figure 5G-I**) showed a preference for F at P3, similar to HLA-B40:01(**Table S15**). These observations are consistent with recognition of P3 by a hydrophobic pocket revealed by structural analysis of ERAP2 (**Figure S14**). Note that cluster 4 was enriched for dependent peptides (**Figure 5I**) and enriched for E at P1 (**Table S15**), a negatively charged amino acid that is resistant to trimming by ERAP2. In summary, immunopeptidome data revealed internal peptide sequence preferences of ERAP2 that shape the ligand repertoire in a HLA class I-specific manner.

## Discussion

In this study, we showed that ERAP2 shapes the HLA-A29 peptidome predominantly by overtrimming peptides carrying susceptible residues at their N-terminus while sparing others carrying a sub-optimal residue at the N-terminal positions. We showed that in the presence of ERAP2 preferred amino acids A, K, and R^44^ are underrepresented, while amino acids F and Y are over-represented at P1, but that these effects on P1 are commonly shared with other class I alleles. Strikingly, we identified that ERAP2 specifically increases the abundance of peptides with a distinct submotif (cluster 2, **Figure 3C**) defined by nonpolar aromatic residues F or Y at P2 that specifically binds to HLA-A29. Replication of these findings in non-related HLA-A29-positive cell lines suggests that these effects of ERAP2 on HLA-A29 are common. Indeed, in known crystal structures of ERAP2 with peptides, the P2 side-chain is accommodated in a very shallow pocket that cannot easily accommodate large residues such as F and Y due to steric clashes with nearby enzyme residues^13^ thus making peptides carrying large hydrophobic bulky residues at P2, poorer substrates (**Figure S14**). Note that we further showed that the effects of ERAP2 on this cluster of peptides is different from ERAP1, which did not show selectivity for this submotif of HLA-A29 (**Figure S6**). This fits with the observation that the pocket in ERAP1 that interacts with P2 provides more space for bulky residues^14^. In fact, using correlation as a metric of the effects of ERAP1 and ERAP2, we show that ERAP1 and ERAP2 show non-redundant effects on the HLA-A29 peptidome (**Figure 3I**), which is in line with genetic studies that revealed that ERAP1 and ERAP2 independently contribute to the disease risk for BU^10^.

Although several studies have shown that ERAPs can trim peptides bound to MHC-I^17,18^, structural studies support that ERAP2 can also trim the N-terminal residues from peptide substrates by first sequestering the entire peptide sequence inside the enzyme’s cavity. There, the peptide substrate interacts with amino acid side chains of the enzyme, which are considered to influence the stability of the interaction and thus the trimming rates of the peptides^13,14^. The exact internal peptide sequence preferences for ERAP2 remain poorly understood. In an attempt to map its relevance to antigen presentation, here we considered the entire peptide sequence to capture the full effects of ERAP2 on the class I immunopeptidomes, and identify functional submotifs which may be missed using traditional single residue or motif analysis. We describe highly reproducible motifs of HLA-A29 and identified that peptides that are destroyed by ERAP2 (i.e., ‘sensitive’ to trimming) showed a strong preference for Leucine at P7 and often are presented by multiple alleles (promiscuity). Although we formally cannot exclude the contribution of residual HLA-A29 molecules in the analysis of HLA-A03:01, data from single-HLA cell lines supported overlap in presented peptides with P7-L (**Figure S13**). Based on the crystal structure of ERAP2^13^, the sidechain P7 can be accommodated within a shallow hydrophobic pocket, which suggests that hydrophobic residues like Leucine would be preferred (**Figure S14**).

Thus, structural analysis indicates that L at P7 is near-optimal for trimming by ERAP2, while bulky residues at P2 (e.g., F) reduce trimming by ERAP2. Therefore, we hypothesize that the increase in peptides with bulky residues at P2 in the presence of ERAP2 is a result of the decreased availability of competing peptides with P7-L due to overtrimming by ERAP2. Importantly, nonpolar aromatic residues F or Y at P3 were associated with peptides that are destroyed in the HLA-B40:01 an HLA-B44:03 peptidome, which is consistent with recognition of P3 by a hydrophobic pocket lined by two other aromatic residues (Tyr892 and Tyr455) that can make favorable pi-stacking interactions with the peptide aromatic side-chain (**Figure S14**). F at P3 was also the most common residue considering all 9-11-mers detected by immunoprecipitation of HLA class I (**Figure S10**). The seemingly contrasting preference of F dependent on the position in the peptide substrate, also suggests that predicting substrate specificity based on widely used fluorogenic aminopeptidase substrates (e.g., R-AMC) or peptide series that vary only the N-terminal residue may obscure the full breadth of substrate specificity for this amino peptidase. We do emphasize that the binding motif of HLA-A29 (and other alleles investigated) can obscure the detection of the full internal sequence preferences of ERAP2, but using the presented peptides as a read-out provides the net effect of any internal sequence preferences on antigen presentation.

We showed that the ERAP-sensitive peptides presented by HLA-A29:02 are promiscuous based on their predicted binding scores for other class I alleles, and their detection in the HLA-A29-negative fraction in mass spectrometry analysis. Since these peptides are also characterized by P1 composition (e.g, A, K, R) that is shared with the other HLA allotypes investigated, it is tempting to speculate that HLA-A29 epitope destruction by ERAP2 is a canonical phenomenon common to class I alleles. This is supported by the observation that HLA class I ligands in general show a depletion for residues A, K, and R at P1 in ERAP2-positive cell lines^34^, which are preferred substrates of ERAP2. High hydrophobicity of T-cell receptor contact residues in presented peptides – in particular a hydrophobic P7 – is associated with immunogenicity^46,47^. Perhaps a canonical function of ERAP2 is to destroy epitopes to lower the immunogenic index of peptide cargo presented. This is supported by observations in cancer immunotherapy, where high ERAP2 expression (the risk haplotype for BU) is a strong prognostic predictor of poor survival in patients receiving checkpoint inhibitor therapy to induce T-cell mediated antitumor immunity^48^. Of interest, the size of P1 of the presented peptide modulates the configuration of position 167 in HLA-A^49^, which was shown to critically influence T cell recognition^47^. F or Y at P1 gives a similar configuration for position 167, which is different from the conformation mediated by K and R at P1 in one study^49^, which suggests that the effects of ERAP2 on P1 may influence T cell receptor recognition.

Given that HLA-A29 is prerequisite for the development of BU, we hypothesize that disease mechanisms associated with antigen presentation are most likely driven by a limited set of epitopes (**Table S8**) because of promiscuity of peptides^45^. ERAP2 destroyed the only S-antigen peptide detected in the HLA-A29 peptidome, which considering high ERAP2 expression is a risk factor for BU, suggests that HLA-A29-mediated presentation of S-antigen fragments is less likely relevant during disease initiation. However, BU patients show in vitro T cell proliferation towards S-antigen.^7^ This makes it tempting to speculate that the retinal S-antigen is more relevant in later stages of the disease via CD4^+^ T cells responses after the blood retina barrier has been breached. This is supported by the common immune reactivity towards S-antigen in patients with clinically distinct phenotypes of uveitis and the lack of response of patient-derived ocular CD8^+^ T cells towards this S-antigen peptide^2,4^. Based on the submotifs of peptides (i.e. cluster 2, **Figure 3C**), we hypothesize that ‘uveitogenic’ HLA-A29-restricted peptides may more likely harbor a F or Y at P2. The importance of P2 is supported by the fact that fine mapping studies of the *MHC* linked BU risk to amino acid positions 62-Leu and 63-Gln of *HLA-A*^50^, which are unique to HLA-A29 and directly interact with P2 of the anchoring peptide. Although the *HLA-C*14:03* allele also showed good binding scores for the ERAP2-dependent peptides with F or Y at P2, *HLA-C* alleles are notoriously low expressed^51^ and the allele frequency of *HLA-C*14:03* is >100 times lower compared to *HLA-A*29:02*. Also, the peptidomes of HLA-A29 and HLA-C14:03 are starkly different (Jaccard similarity index ±1% using peptidome data from Sarkizova and associates^31^) and T cells recognizing the same peptide in a different HLA molecule may not show immune reactivity. Although the overall binding motif of peptides bound to HLA-A29:02 is similar to other alleles (HLA-A1 family members, such as HLA-A01 and HLA-A:30:02^31^), our unbiased submotif analysis had the resolution to discover functional differences not apparent when considering the overall motif of HLA allotypes. The peptide motif of cluster 2 in particular affected by ERAP2 and presented by HLA-A29 (**Figure 3C**) did not bind well to the other alleles (**Figure 3J**) and supports that HLA-A29 has unique features in antigen presentation, and is in line with the fact that no other HLA alleles are genetically associated with BU^9^. Note that ‘just’ P2-F and P2-Y (so without the C-terminal position Y) is not uncommon in the immunopeptidomes of other HLA-A allotypes, such as HLA-A24. We previously showed that peptides with the P2-F/Y+P**Ω**-Y motif are infrequent on functionally similar HLA allotypes or that these HLA allotypes display very low similarity in the immunopeptidome composition with HLA-A29^52^. This supports that ERAP2 can influence the presentation of peptide that are specific for HLA-A29

Regardless, we show that the amino acid sequence of retina-expressed genes contains peptides with the motif of cluster 2, which supports that ERAP2-mediated HLA-A29-restricted presentation of ocular epitopes could be a key disease mechanism for BU. We hypothesize that ERAP2 facilitates higher expression of HLA-A29-specific peptides derived from proteins related to melanocyte biology – specifically expressed in the ocular retina or choroid. Of course, functional experiments of antigen presentation in the eye and tetramer-analysis of T cell immunity to these putative epitopes is warranted. It is, however, of interest that among the predicted epitopes we found peptides derived from key factors in melanocyte biology. A hallmark feature of BU is the progressive loss of stromal melanocytes in the choroid corresponding to the characteristic cream-colored birdshot fundus lesions^2,8,53,54^, and BU has been associated with melanoma^5,6,55^.

Previous HLA peptidomic studies of ERAPs are based on single-HLA or long-established cell lines which after years of continuous cultivation are notorious for their profound chromosomal aberrations reported to also affect *ERAP* and *HLA* genes^44,56–58^. In addition, these studies have been conducted with label-free approaches using independent experimental runs, which makes accurate quantification of effects of ERAPs on the immunopeptidome more challenging. To study ERAPs in a physiologically more relevant environment, we exploited MS analysis using newly-established patient-derived cell lines and SILAC labeling to address several potential sources of ambiguity that are non-trivial to resolve with *in silico* methods, including often unaccounted genetic variability (i.e., polymorphisms) in comparing different cell lines or quantitative error caused by the individual analysis of to be compared conditions. Regardless, the results in this study can also be influenced by several factors. Although abundant peptides are more likely to be sufficiently detected in individual elutions (~90% of peptides were reported before), less abundant peptides might be missed. This means that additional undiscovered effects of ERAP2 on the peptidomes investigated could be present. For example, we limited our labeling and analysis to peptides that contain F and/or Y for SILAC labeling, which obscured our capability to cover the majority of the HLA-B40:01 peptidome or potential uncharted domains of the peptidomes of the other alleles.

In conclusion, we show that ERAP2 significantly influences the immunopeptidome across the cellular HLA class I allotypes and ERAP2 increases the expression of a peptide submotif highly selective for HLA-A29. We have narrowed down the potential sequences for ocular-derived antigenic peptides based on the selective effect of ERAP2 on the peptide cargo of HLA-A29 in the pathogenesis of Birdshot Uveitis.

## Acknowledgements

We express gratitude for constructive input from prof. Debbie van Baarle.

## Supplemental Info for

### Supplemental Methods and Info

#### Lentiviral vector production

HEK-293T cells were seeded into 10 cm dishes (2×10^6^ cells/dish) and cultured in Dulbecco’s Modified Eagle Medium (DMEM, Thermo Fisher Scientific). The next day, 293T cells were co-transfected with 2 μg transfer vector (Lenti ORF clone of Human S-antigen mGFP tagged, RC220057L2 from Origene) and components of 2nd generation packaging vectors: 8.33 μg psPAX2 packaging vector and 2.77 μg pMD2.G envelope vector at a ratio of 4:1. Transfection was done in serum-free DMEM using Lipofectamine 2000 (Thermo Fisher Scientific) according to manufacturer’s instructions. Medium was replaced with 10 mL DMEM supplemented with 10% FBS and incubated at 37°C, 5% CO_2_ after 24 hours. The conditioned medium containing lentiviral particles was collected 48 hours after transfection and an additional 10 mL of fresh culture medium was added to the cells. After 12 hours, harvested supernatants were combined and cleared by centrifugation at 1500 rpm for 5 minutes at 4°C then passed through a 0.45 μm filter. Lentiviral supernatants were concentrated using ultracentrifugation with a Beckman Coulter Optima centrifuge using a SW32Ti rotor. Filtered supernatant was added to 38.5 mL Ultra-Clear tubes (Beckman Coulter). Centrifugation was performed for 120 minutes at 32,000 rpm. Supernatant was completely removed and virus pellets were resuspended in 1 mL RPMI (containing 10% FBS and 1% penicillin/streptomycin) and stored at −80°C.

#### Lentiviral transduction of S-antigen in EBV-LCL

To obtain stable cell lines overexpressing S-antigen, EBV-LCLs were transduced with the concentrated lentiviral supernatants. To transduce EBV-LCLs, 1×10^6^ cells were seeded in a 24-wells plate with the lentivirus and a final polybrene concentration of 6 μg/mL. After 24 hours, the medium was replaced and the cells were cultured for another 3 days, without exceeding a cell concentration of 1.5 × 10^6^ cells/mL. Transduction efficiency was monitored by fluorescent light microscopy. GFP-positive EBV-LCLs were sorted using the BD FACSAria^™^ III sorter and S-antigen expression levels were detected by western blot. Western blot analysis to detect the fusion protein S-antigen-GFP was done as described in the Method section. Figure 1 below shows the high protein expression of S-antigen and Tubulin detected from the same cell lysate samples of the patient-derived LCL, but run on two separate blots in parallel because the proteins are detected in close proximity on the blot.

**Figure 1.**
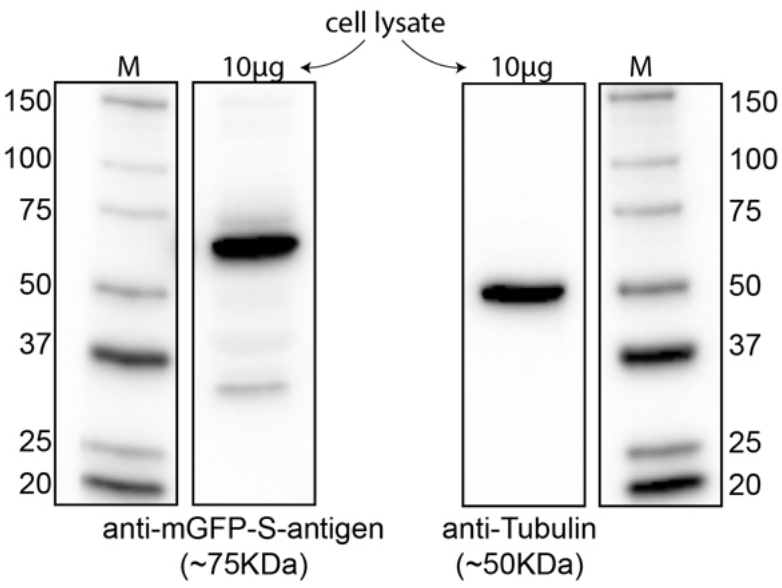
Westernblot analysis of retinal S-antigen after transduction of the patient-derived LCLs.

#### Differential expression analysis of peptides using limma

For differential expression analysis we used the workflow from Kammers *et al*., 2015 available at http://www.biostat.jhsph.edu/~kkammers/software/eupa/R_guide.html. Their method exploits the R package *limma* for shrinking a peptide’s sample variance towards a pooled estimate that boosts power for stable detection of (truly) significant changes in small proteomic data sets. Peptide data were preprocessed using the *read.peptides()* function, which excludes peptides with missing values (i.e., not detected in either the light or heavy channel). We computed dummy variables for the “*Isolation.Interference”, “Quan.Usage”, “Quan.info*” variables, because quality control of the input data was completed as described in the main manuscript. The peptide sequence was used as the “*Protein.Group.Accessions*” variable. Overlapping peptide data from the biological replicates were independently normalized using the *quantify.proteins()* function. Following the workflow of Kammers et al., we used peptides (with a Mascot Percolator q<0.01 in all analyses) detected in both biological replicates (i.e., peptides unique to one of the conditions are left out for normalization and statistical analysis). For example, for peptides detected by DK1G8 (anti-HLA-A29) with a *HLAthena* binding score [MSi]>0.6 for HLA-A*29:02 a total of 1330 peptides were detected in both channels, while 41 peptides in either the light or heavy channel (with consistent detection in the same channel in both experiments) and were not considered for statistical analyses. We blocked for batch effect (two independent experiments) in *limma* by including them in the design matrix. HLA-A29 peptidome analysis considering also peptides detected in either the heavy or light channels is provided in **Figure S3**. Here, we used dummy variables for the moderate q-value (set to 1 × 10^-6^) and log2FC (log_2_FC= −6.6 for peptides only detected in the ERAP2 KO-cell line and log2FC=6.6 for peptides detected only in the ERAP2 WT-cell line), because these parameters were only used to subset peptides unique to either of the conditions (using moderate q<0.01 as a threshold) together with the differentially expressed peptides detected in both channels. Also, although Mascot Percolator exploits a number of relevant peptide features and has been shown to be superior in accurate peptide identification compared to previous Mascot scoring based on one metric (Borsch et al.,2009), we also conducted this analysis of the HLA-A29 peptidome using the percolator q-value in conjunction with the Mascot ions score >30, which showed similar effects for ERAP2 at the submotif level as the analysis using the percolator q-value (see **Figure S3**).

### Supplemental Figures

**Supplemental Figure S1.**
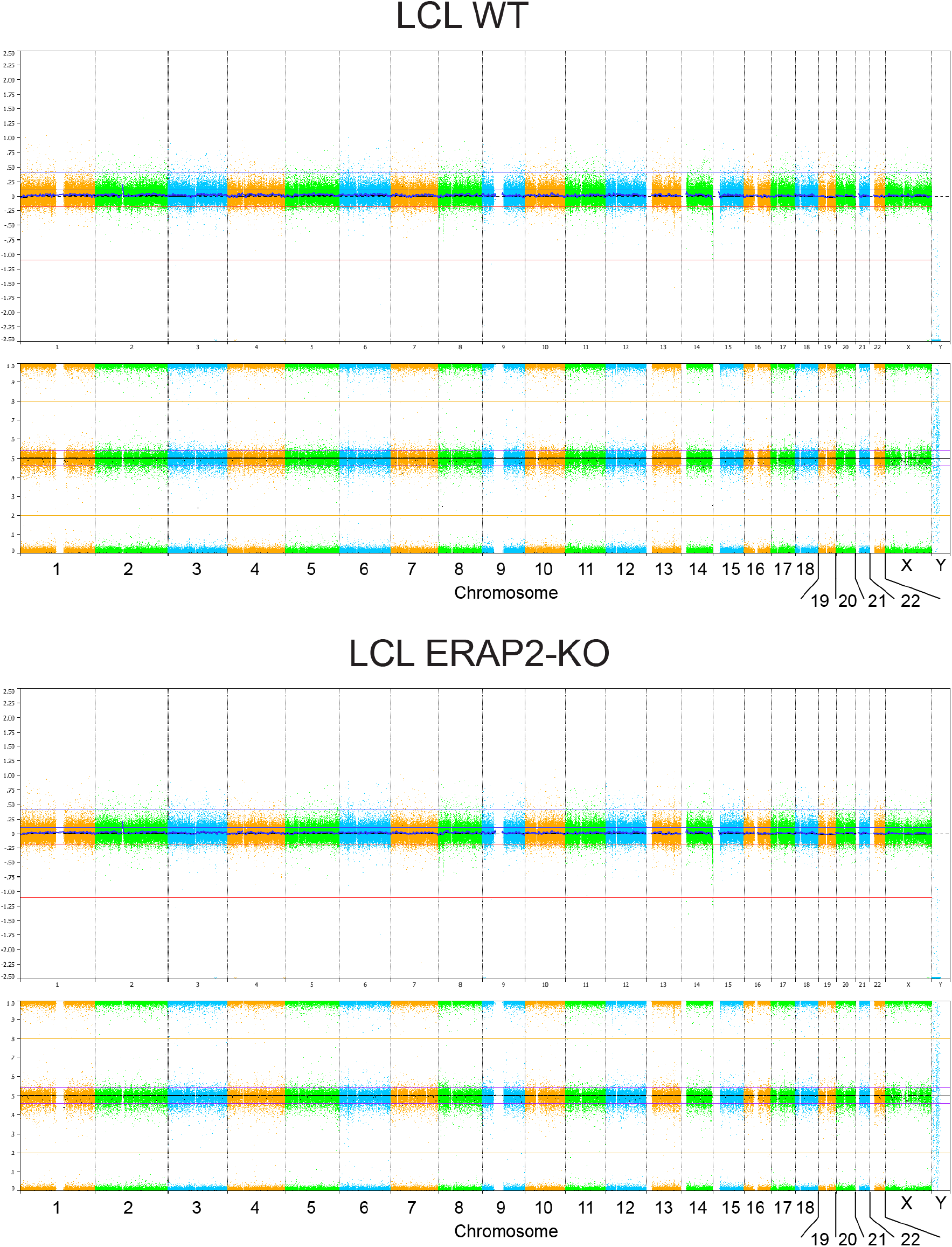
Whole genome analysis using SNP arrays on unedited (LCL Wildtype, WT) and edited (ERAP2-KO) cell lines used in this study. SNPs were detected by the Infinium Human CytoSNP-850K v1.1 BeadChip (Illumina, San Diego, CA, USA) and show highly consistent genomes. The panels show the array results for the whole. On the X-axis the chromosomes and chromosomal region are indicated. The upper Y-axis shows the Log2 R ratio and the lower Y-axis indicates the B allele frequency for each SNP.

**Supplemental Figure S2.**
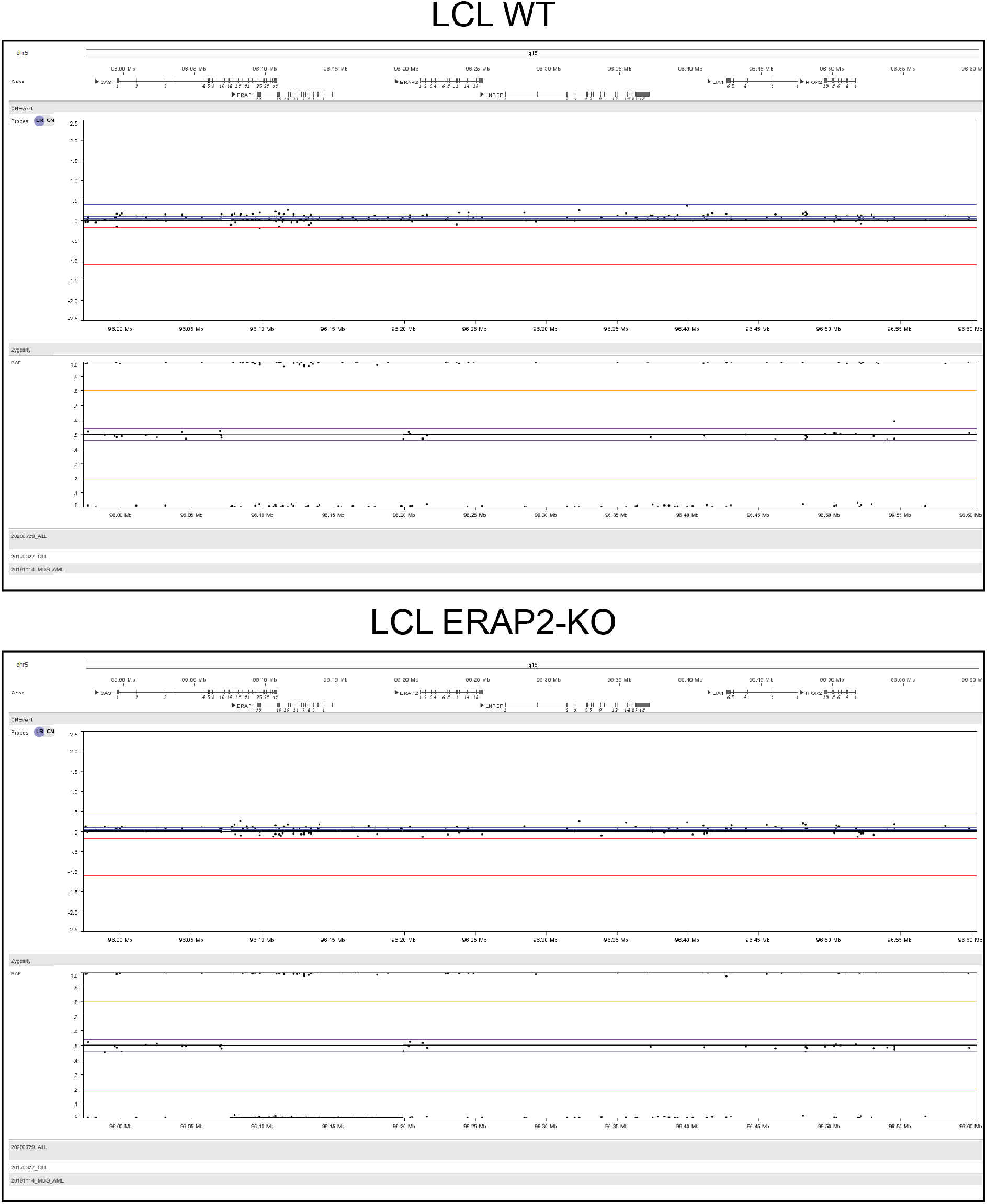
Whole genome analysis using SNP arrays on unedited (LCL Wildtype, WT) and editec (ERAP2-KO) cell line using the Infinium Human CytoSNP-850K v1.1 BeadChip (Illumina, San Diego, CA, USA), similar to **Supplemental Figure S1**, but here the panels show the array results for the region near *5q15* including *ERAP1, ERAP2*, and *LNPEP*. The SNP probes are indicated by black dots. The upper Y-axis shows the Log2 R ratio for the probes and the lower Y-axis indicates the B allele frequency for each SNP (BAF).

**Supplemental Figure S3.**
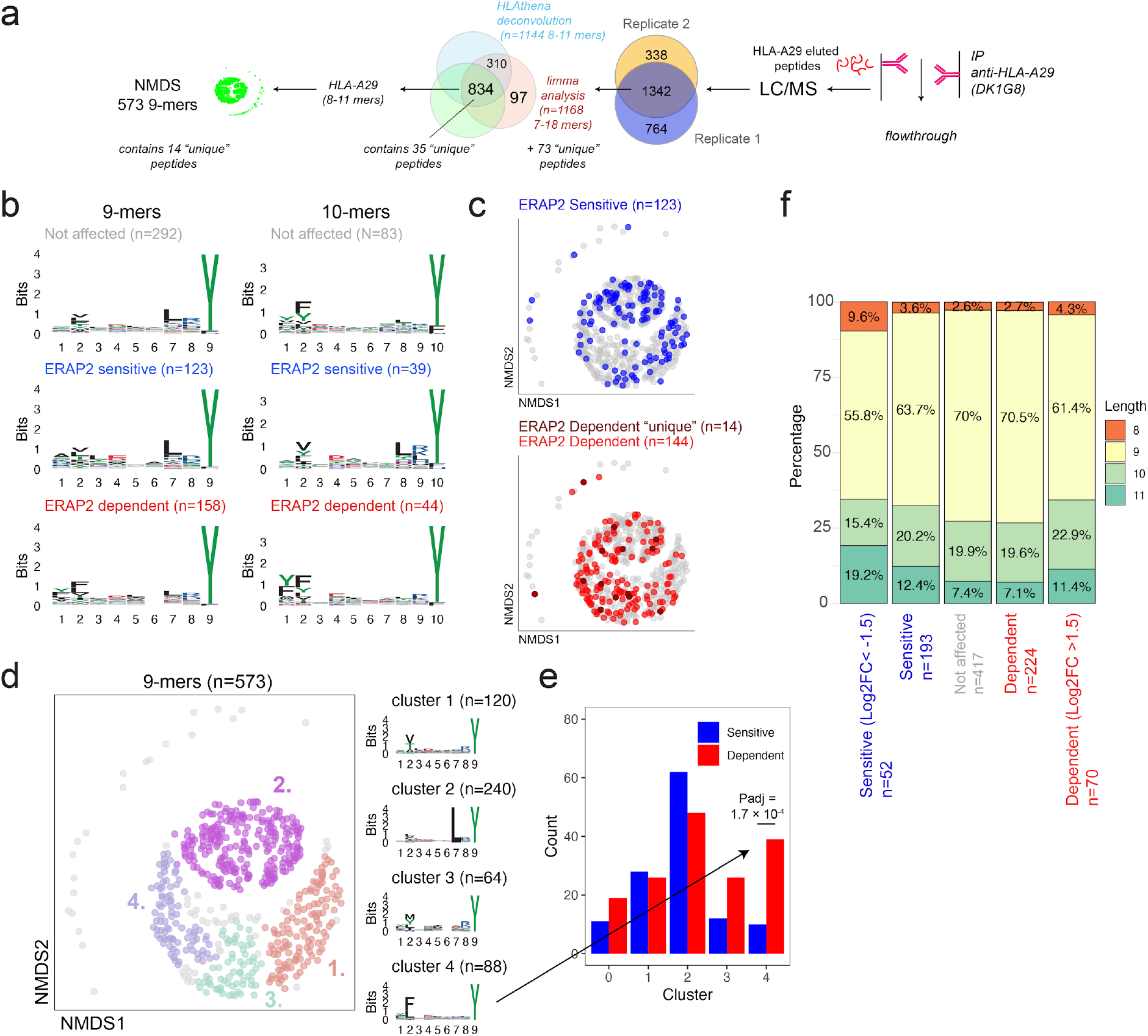
HLA-A29 peptidome data analysis including peptides unique to either ERAP2-KO or ERAP2-WT cells. **a**) A total of 1342 peptides overlapping between the two biological replicates with percolator q<0.01 and Mascot Ions score >30, were filtered according to the steps indicated. Note that after the limma analysis, the 73 “unique” peptides detected in either the heavy or light labeled conditions (with consistent detection in the same channel in both experiments) were added to the dataset before deconvolution with *HLAthena* to filter for HLA-A29 ligands. **b**) The sequence logos for 9-mers and 10-mers in this dataset. ERAP2-sensitive peptides are peptides that decrease in amount in the presence of ERAP2 and ERAP2-dependent peptides increase in amount in the presence of ERAP2. **c**) Nonmetric multidimensional scaling of 573 9-mers in this dataset. The ERAP2-sensitive and ERAP2-dependent peptides are indicated in blue and red, respectively. Peptides uniquely identified in the ERAP2 WT-condition are shown in dark red (n=14). **d**) Four clusters were estimated (eps parameter for DBSCAN, using k=5) using the elbow method. The sequence logos for each cluster are indicated on the right. **e**) Comparison of the number of ERAP2-sensitive and ERAP2-dependent peptides in each peptide cluster identified in *b. Padj* = bonferroni corrected (n=clusters) *P* values from X^2^ tests. All other comparisons were Padj>0.05. **f**) The percentage of 8-11-mers in peptides sets of this dataset. This analysis shows length dependent effects seen for ERAP2 in an hypoactive ERAP1 background.

**Supplemental Figure S4.**
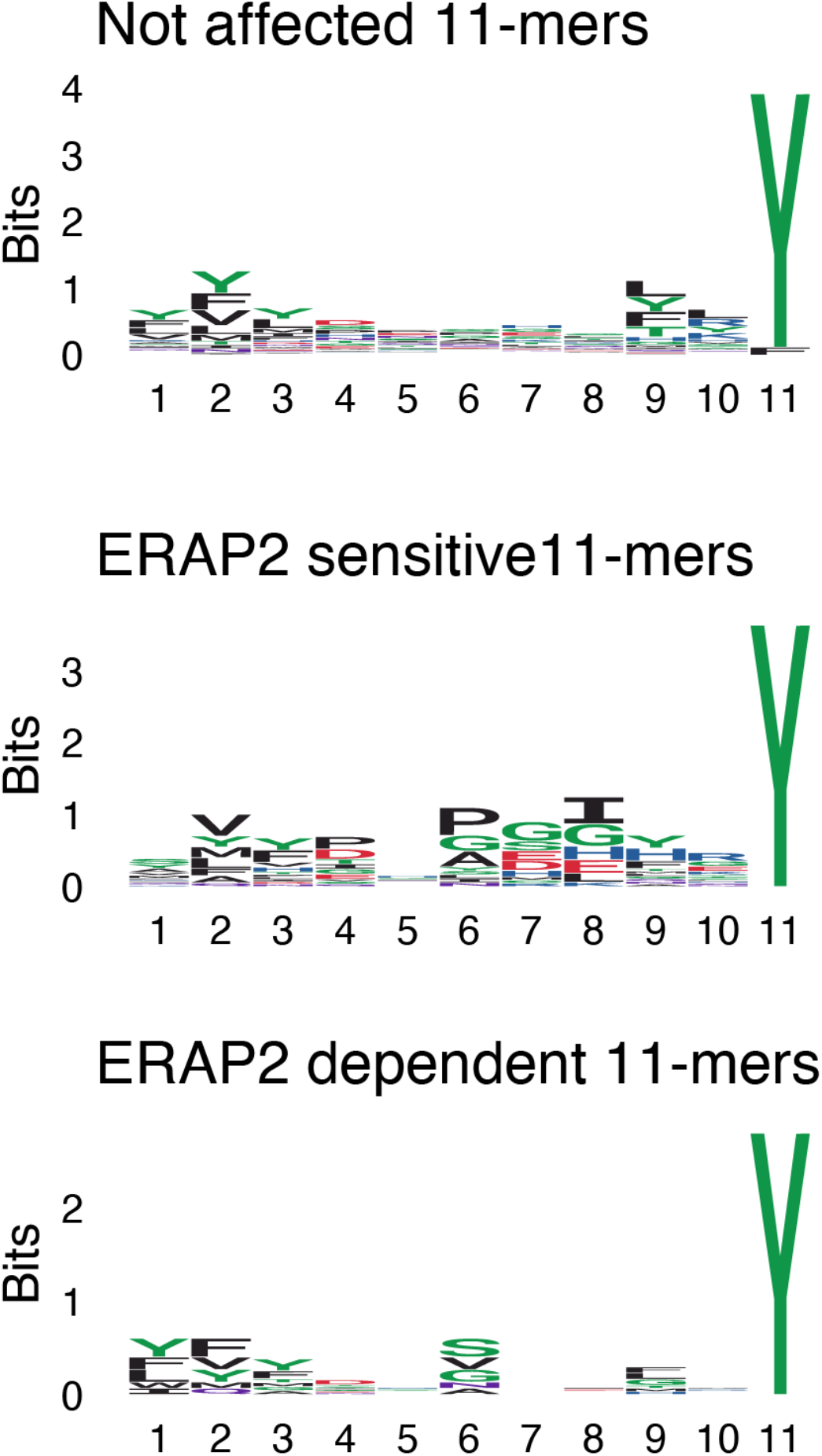
The sequence logos for non-redundant 11-mers from HLA-A29. Peptides that decrease in the presence of ERAP2 are termed ERAP2-sensitive, peptides that increased in relative amounts are termed ERAP2-dependent. Peptides that did not change in relative amounts in the presence of ERAP2 are termed ‘not affected’.

**Supplemental Figure S5.**
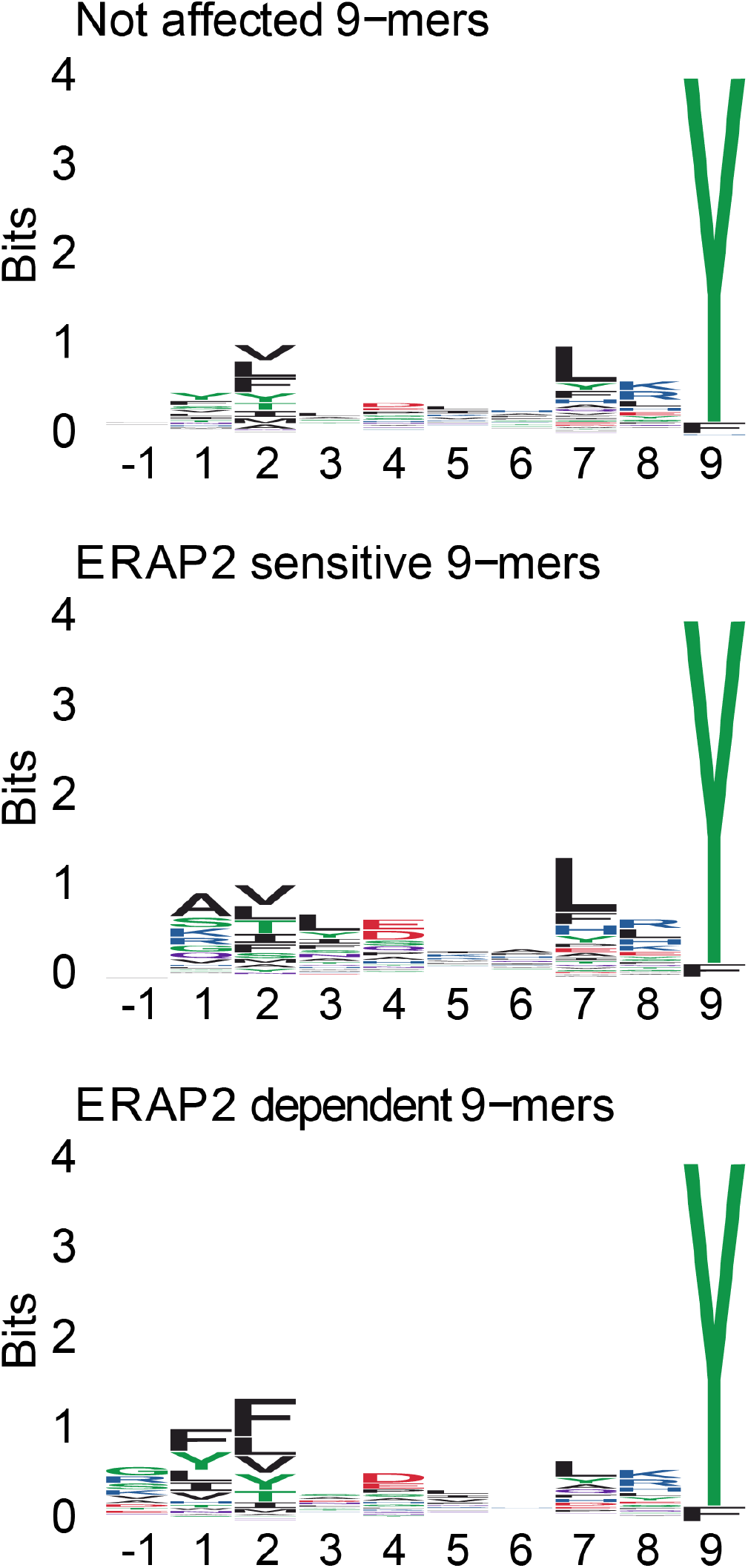
The sequence logos for 948 non-redundant 9-mers and their designated P-1 derived from the amino acid sequence of the putative proteins. Peptides that decrease in the presence of ERAP2 are termed ERAP2-sensitive, peptides that increase in abundance are termed ERAP2-dependent. Peptides that did not change in abundance in the presence of ERAP2 are termed ‘not affected’.

**Supplemental Figure S6.**
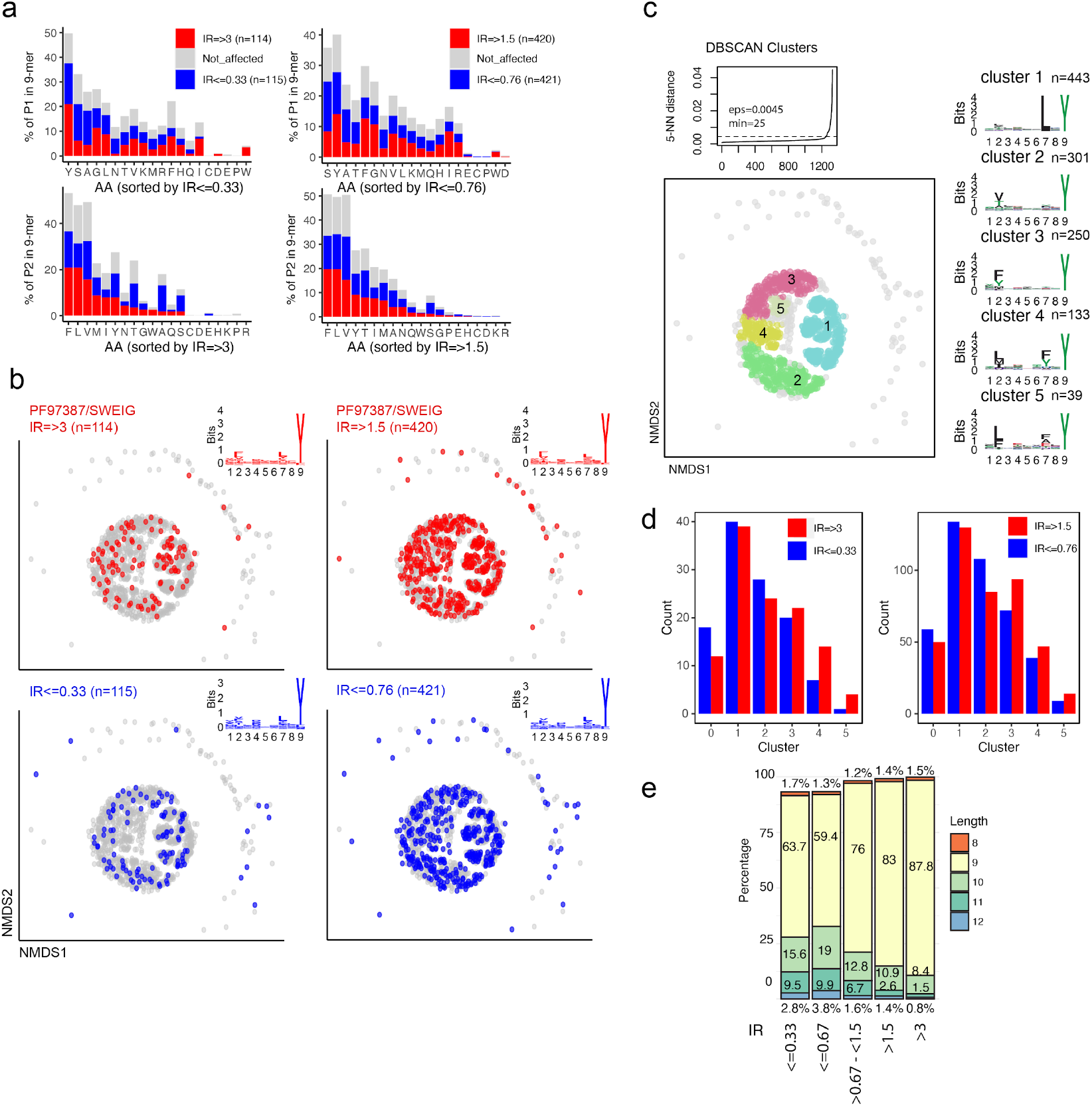
Non-metric multidimensional scaling of the 1329 shared 9-mers eluted from the HLA-A29-positive cell lines PF97387 (ERAP1 high expression/activity) and SWEIG (ERAP1 low expression/activity). The 1329 9-mers were filtered (removed peptides with value 0 in any of the 3 replicates from PF97387 or SWEIG) from a total of 5584 (3828 9-mers) peptides from *Alvarez-Navarro et al., 2015*. In this study, the normalized intensity ratio (PF97387/SWEIG) of each peptide in the two cell lines was used to infer the relative abundance of each peptide, which we adapted to assign peptides as ERAP2-dependent (IR ≥ 1.5 or IR ≥ 3) or ERAP2-sensitive (IR ≤ 0.76 or IR ≤ 0.33). We used IR≤ 0.76 (instead of 0.67) compared to IR ≥ 1.5 so the peptide datasets would be of equal size. **a**) Comparison of amino acid proportion at P1 and P2 of 9-mers (in percentage for each group of peptides) between peptides that decrease in abundance (in blue) in the presence of ERAP1 or that increase in abundance (red), compared to peptides not affected by ERAP1 cells (in grey). All comparisons were not significant; *Padj>0.05*. **b**) Non-metric multidimensional scaling of the 1329 9-mers **c**) Five clusters were estimated (eps parameter for DBSCAN, using k=5, based on Figure 3C) using the elbow method. The sequence logos for each cluster are indicated on the right. **d**) Comparison of the number of ERAP1-dependent (IR ≥ 1.5 or IR ≥ 3) and ERAP1-sensitive peptides (IR ≤ 0.76 or IR ≤ 0.33) in each peptide cluster identified in *b*. *Padj* = bonferroni corrected (n=clusters) X^2^ tests. The difference between the count of sensitive and dependent peptides in each cluster was not significant or *Padj>0.05*. **e**) The percentage of 9-mers and 10-mers in peptides sets using different cut-offs for the intensity ratio (IR). This analysis confirms the length effects seen for ERAP1 as reported by *Alvarez-Navarro et al., 2015* in these cell lines (which are ERAP2-deficient).

**Supplemental Figure S7.**
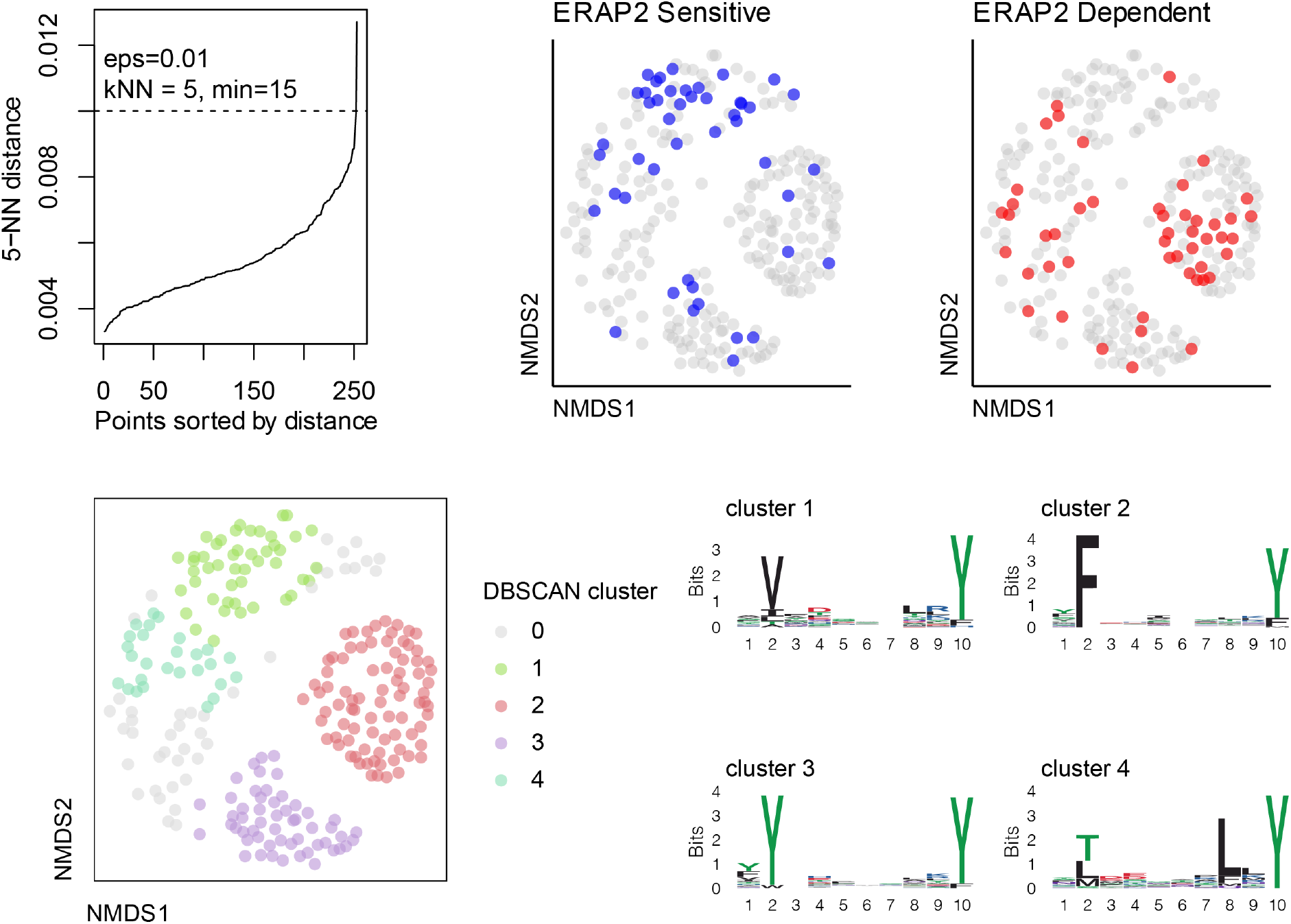
Non-metric multidimensional scaling plot of 235 10-mers eluted from HLA-A29:02. Differentially expressed peptides are indicated in blue (ERAP2 sensitive that decrease in abundance in the presence of ERAP2) and red (ERAP2-dependent peptides that increase in abundance in the presence of ERAP2). A total of four clusters were identified and the sequence logos for each cluster are indicated. Cluster 0 indicates the unassigned peptides. 10-mer peptides of cluster 2 also show the P2-F motif for HLA-A29 and contains enrichment for ERAP2-dependent peptides compared to ERAP2-sensitive peptides, which reflects the results from 9-mers in **Figure 3**.

**Supplemental Figure S8.**
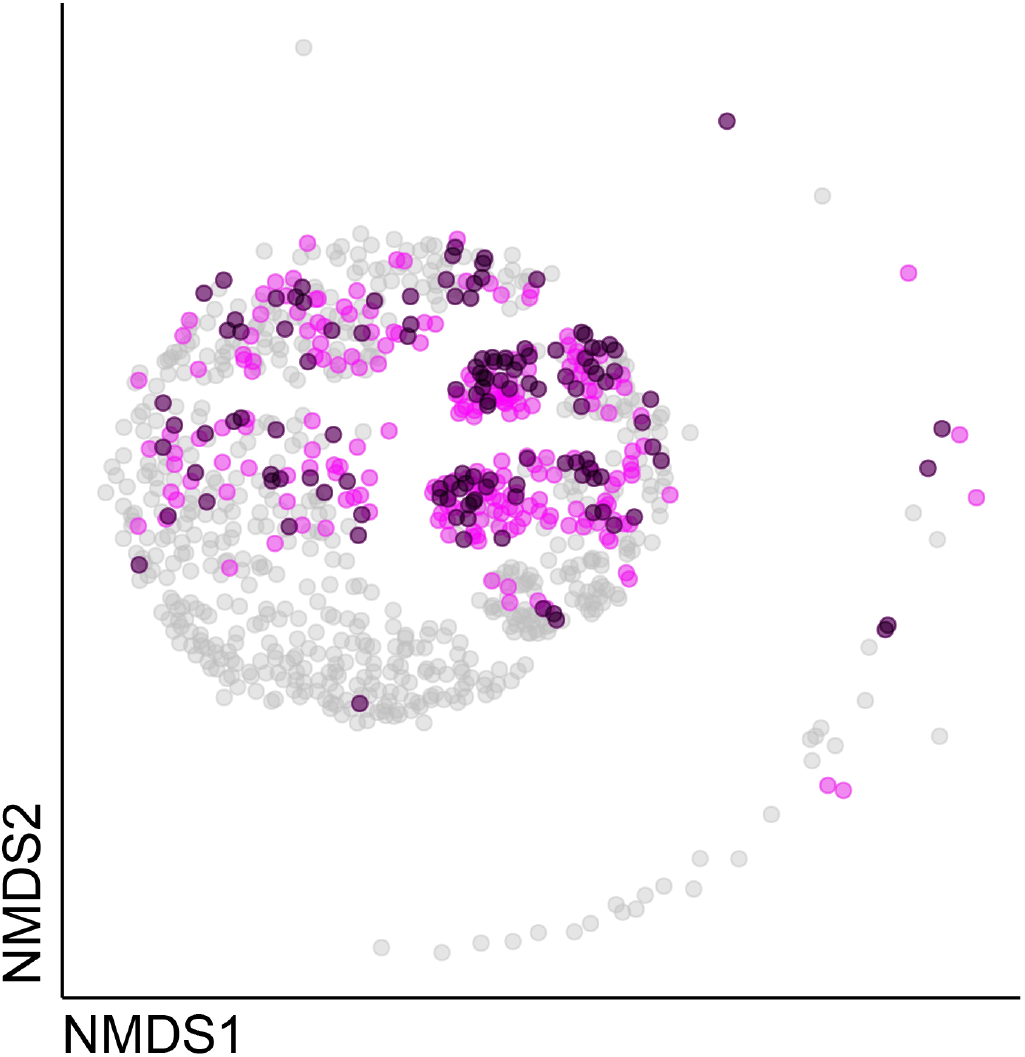
Non-metric multidimensional scaling plot of 948 9-mers eluted from HLA-A29:02 in this study. Peptides with a binding score MSi>0.6 for HLA-A03:01 from *HLAthena* (https://HLAthena.tools) are highlighted in magenta. Peptides with a binding score MSi>0.6 for HLA-A03:01 that are differentially expressed (moderate q<0.01) are indicated in black.

**Supplemental Figure S9.**
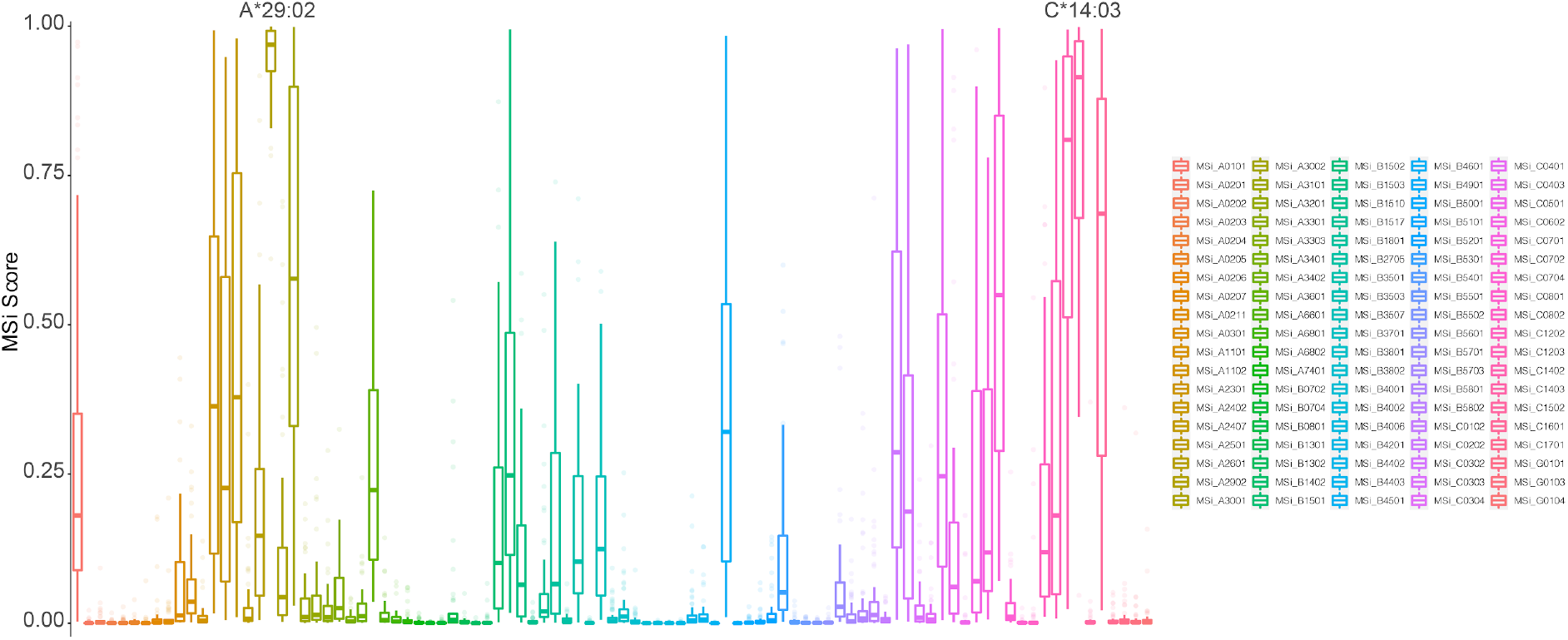
Predicted binding scores (in MSi from HLAthena) for the 53 ERAP2-dependent peptides in cluster 2 (**Figure 3C**) across 95 HLA alleles (selection of alleles tested based on Sarkizova *et al., 2020*).

**Supplemental Figure S10.**
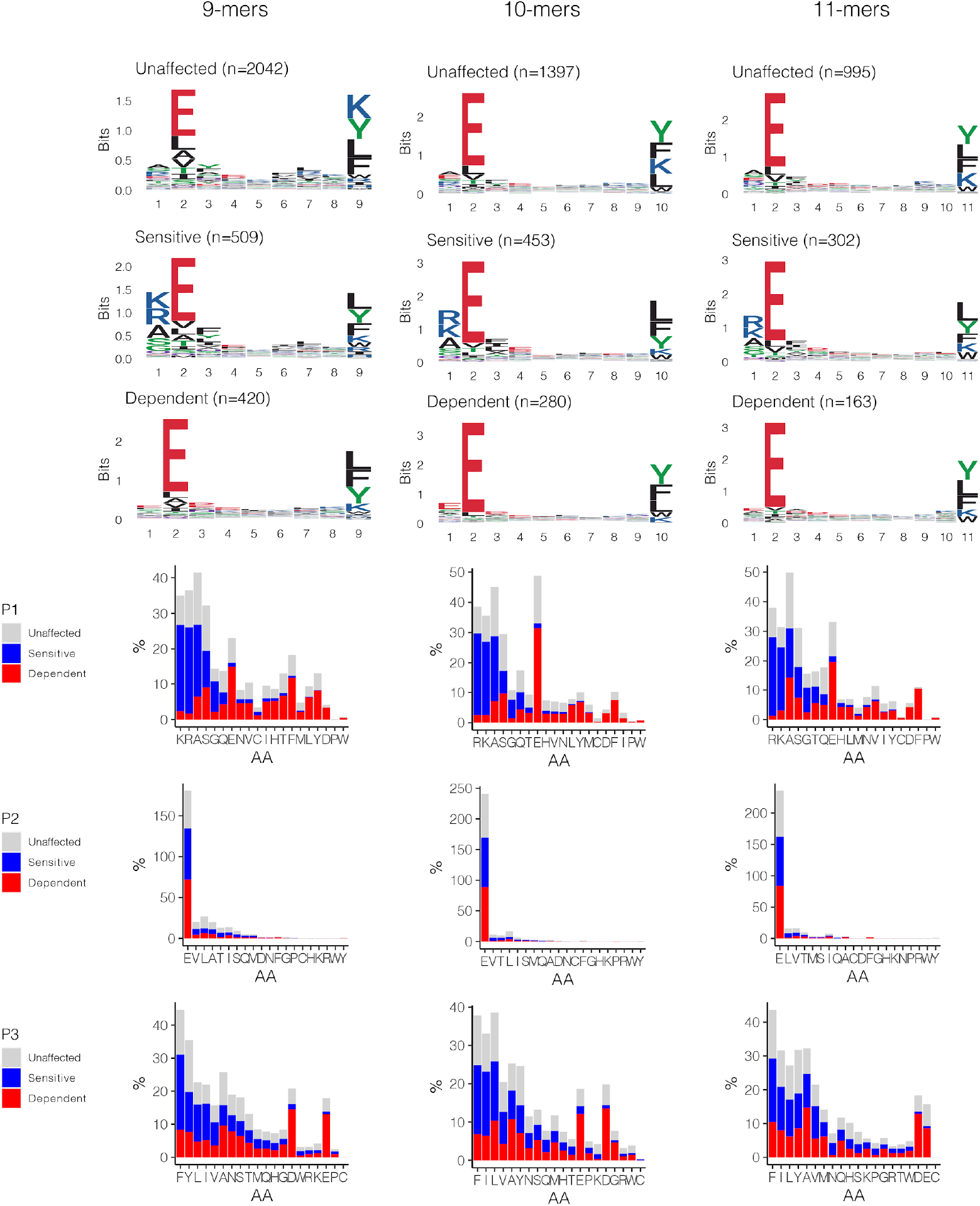
The effect of ERAP2 on the HLA class I peptidome. Sequence motifs depict specific amino acid preferences for 9-, 10-, and 11-mers were generated from a non-redundant list of peptides from HLA class I (W6/32). Comparison of amino acid proportion at P1, P2, and P3 of (in percentage for each group of peptides) between peptides that decrease in abundance (‘sensitive’), peptides that increase in abundance (‘dependent’ peptides) compared to peptides not affected in ERAP2-WT cells (in grey).

**Supplemental Figure S11.**
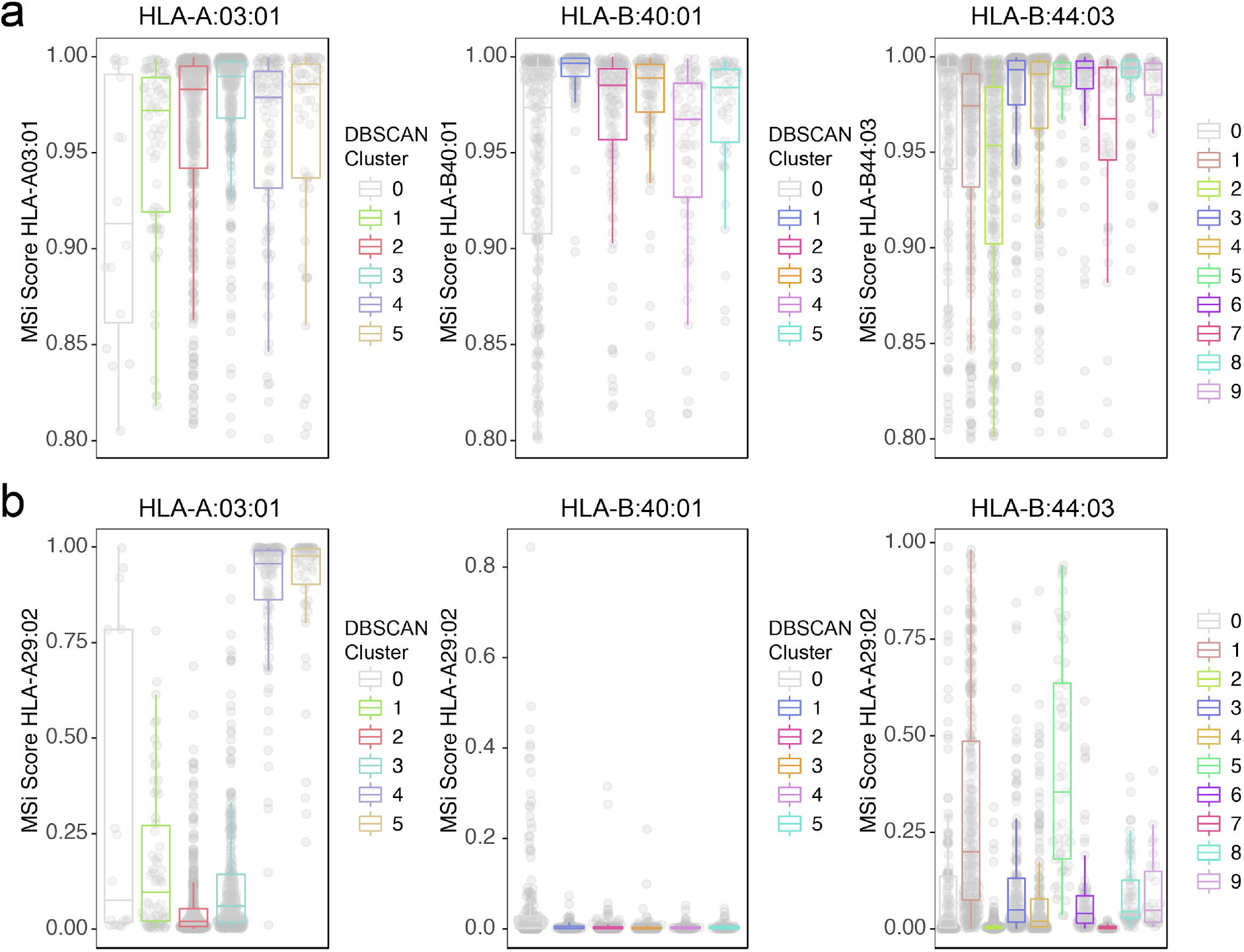
Peptide bindings scores from *HLAthena* (HLAthena.tools) for peptide clusters from **Figure 4**. **a**) The binding score (MSi) for 9-mers with a MSi>0.8 used for the non-metric multidimensional scaling of *HLA-A*03:01, HLA-B*40:01*, and *HLA-B*44:03*. The binding score ranges from 0 (low) to 1 (high). Clusters identified by DBSCAN are indicated and color-coded. **b**) The binding score for *HLA-A*29:02* (MSi) for the same 9-mers and clusters as shown in *a*.

**Supplemental Figure S12.**
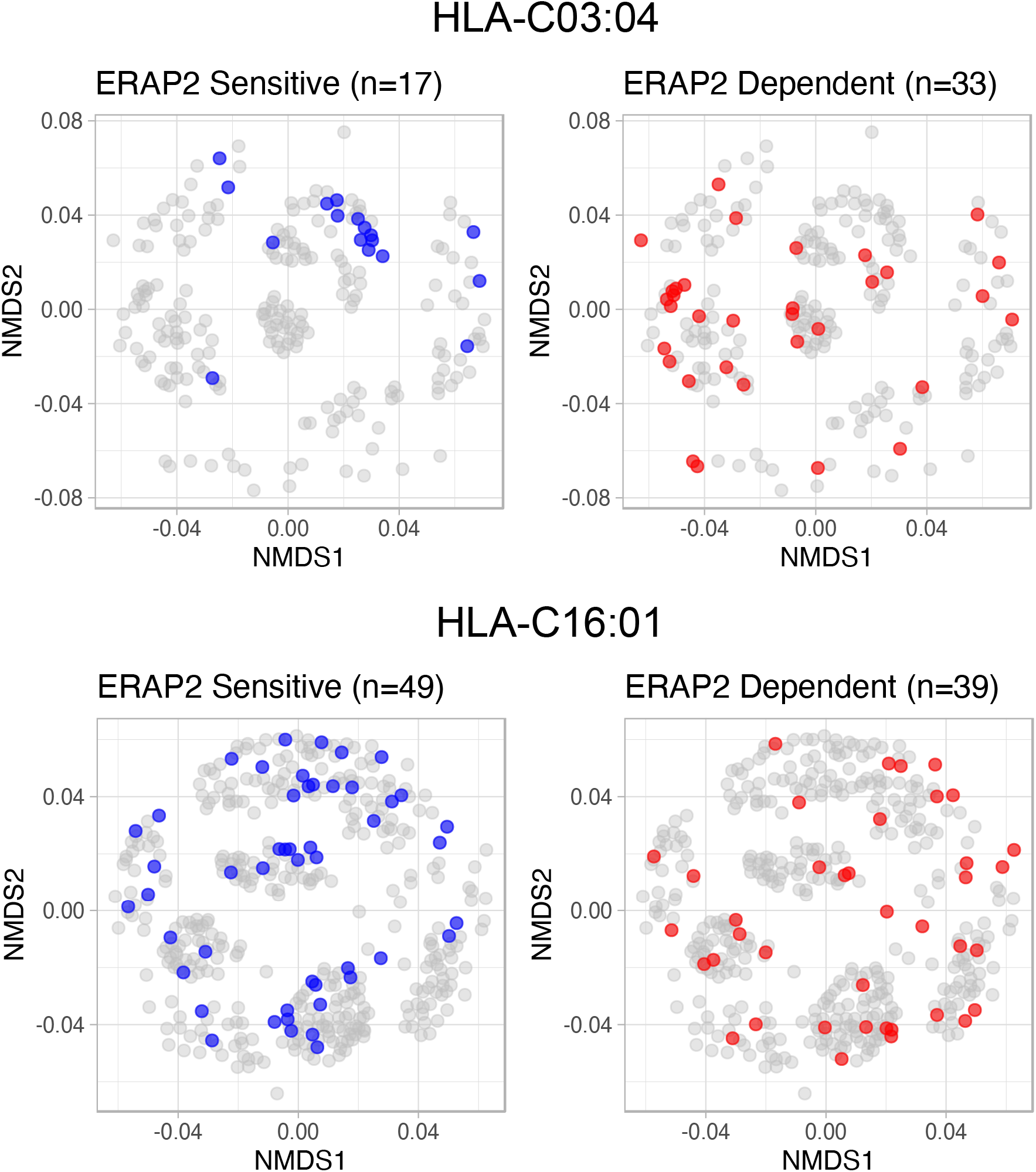
Non-metric multidimensional scaling plot of 221 and 411 9-mers from HLA-C03:04 and HLA-C16:01. Differentially expressed peptides are indicated in blue (ERAP2 sensitive that decrease in abundance in the presence of ERAP2) and red (ERAP2-dependent peptides that increase in abundance in the presence of ERAP2).

**Supplemental Figure S13.**
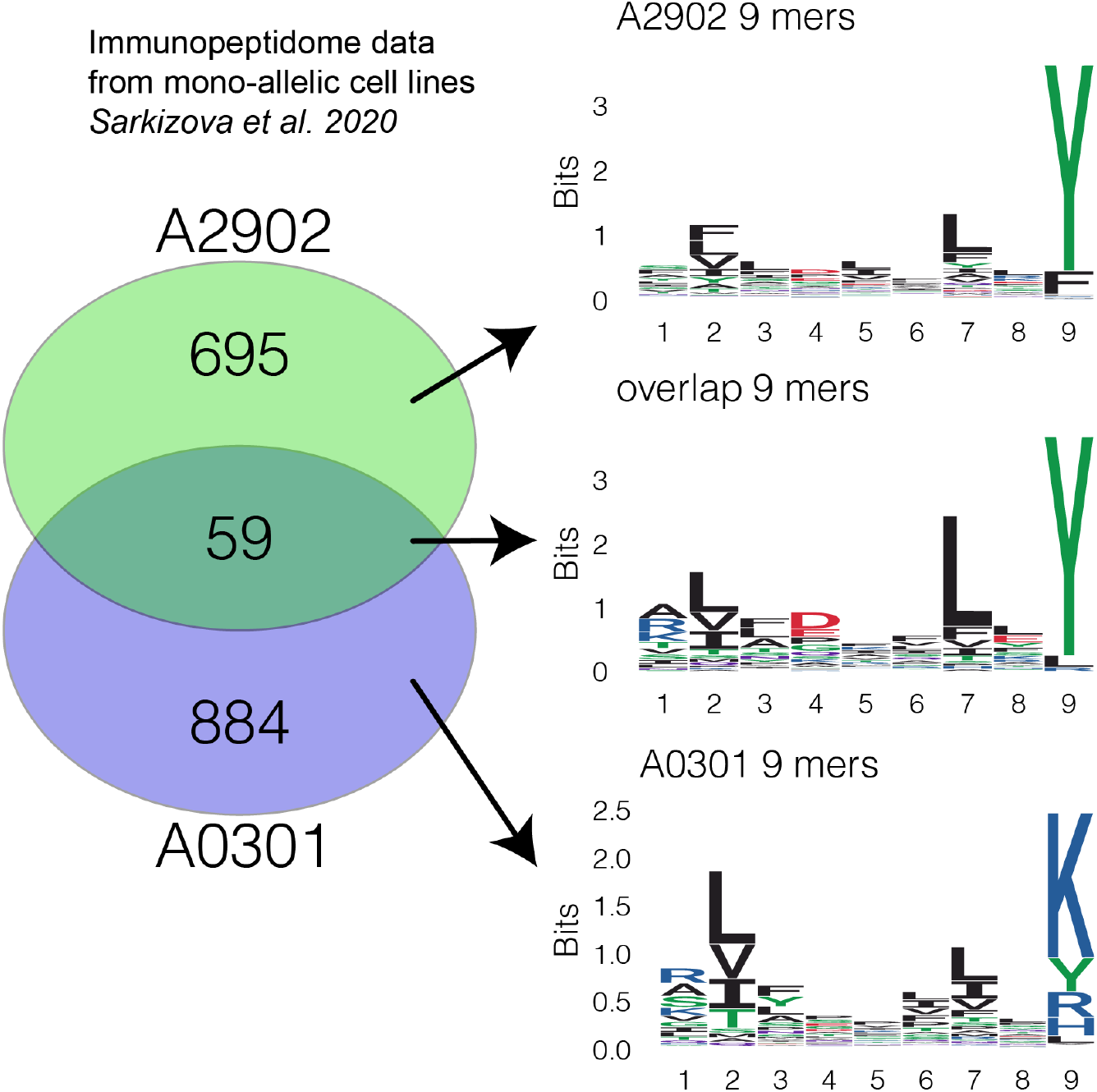
Venn diagram of 9-mers presented by monoallelic cell lines expressing only HLA-A29:02 or only HLA-A03:01 from *Sarkizova et al., 2020* A total of 59 9-mers were detected in both datasets. The sequence logos for peptides uniquely observed in HLA-A29, overlapping peptides found in both monoallelic datasets, and peptides uniquely observed in HLA-A03 are indicated on the right.

**Supplemental Figure S14.**
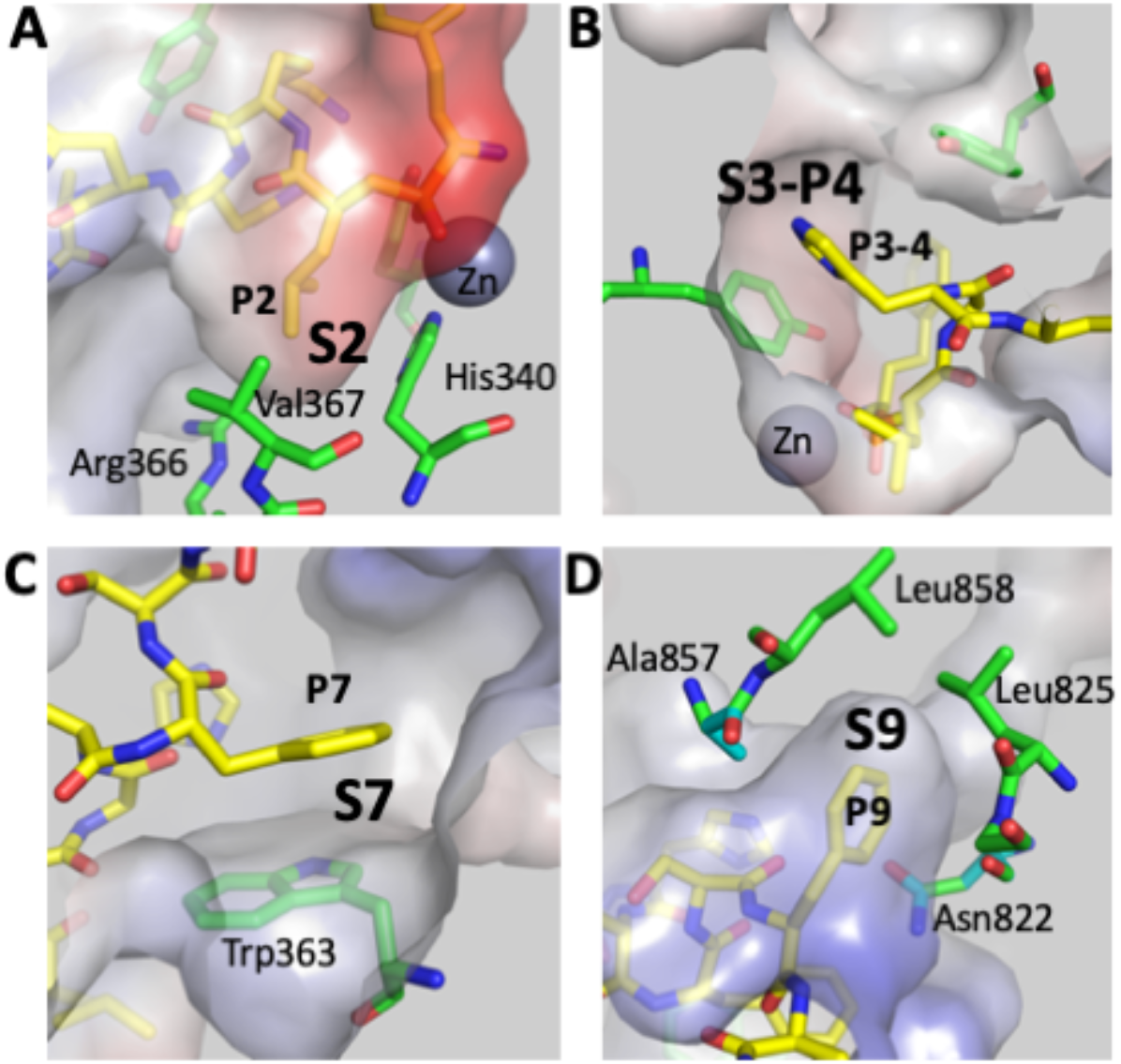
Putative specificity pockets of ERAP2 that help explain observed sequence motifs. ERAP2 (from PDB code *5AB0*) is shown in surface representation colored by electrostatic potential (red=negative, white=neutral, blue=negative). Peptide analogue DG025 that was crystallized bound onto ERAP2 is shown in yellow sticks (carbon=yellow, oxygen=red, nitrogen=blue). Nearby ERAP2 residues that help form indicated specificity pockets are shown in green sticks. Specificity pockets are indicated as **a**) S2, **b**) S3-P4, **c**) S7 and **d**) S9. Peptide residue side-chains that are accommodated in the pockets are indicated as P2, P3-4, P7 and P9.

## References

1. Minos E, Barry RJ, Southworth S, Folkard A, Murray PI, Duker JS, et al. Birdshot chorioretinopathy: Current knowledge and new concepts in pathophysiology, diagnosis, monitoring and treatment. Orphanet J Rare Dis. (2016) 11(1):1–17. 10.1186/s13023-016-0429-8

2. Kuiper J, Rothova A, de Boer J, Radstake T. The immunopathogenesis of birdshot chorioretinopathy; a bird of many feathers. Prog Retin Eye Res. (2015) 44:99–110. 10.1016/j.preteyeres.2014.11.003

3. Kuiper JJW, Mutis T, de Jager W, de Groot-Mijnes JDF, Rothova A. Intraocular interleukin-17 and proinflammatory cytokines in HLA-A29-associated birdshot chorioretinopathy. Am J Ophthalmol. (2011) 152(2):177–182.e1. 10.1016/j.ajo.2011.01.031

4. Kuiper JJW, Rothova A, Schellekens PAW, Ossewaarde-van Norel A, Bloem AC, Mutis T. Detection of choroid-and retina-antigen reactive CD8+ and CD4+ T lymphocytes in the vitreous fluid of patients with birdshot chorioretinopathy. Hum Immunol. (2014) 75(6):570–577. 10.1016/j.humimm.2014.02.012

5. Pulido JS, Canal I, Salomão D, Kravitz D, Bradley E, Vile R. Histological findings of birdshot chorioretinopathy in an eye with ciliochoroidal melanoma. Eye. (2012) 26(6):862–865. 10.1038/eye.2012.10

6. Gaudio PA, Kaye DB, Crawford JB. Histopathology of birdshot retinochoroidopathy. Br J Ophthalmol. (2002) 86(12):1439–1441. 10.1136/bjo.86.12.1439

7. Nussenblatt RB, Mittal KK, Ryan S, Richard Green W, Edward Maumenee A. Birdshot Retinochoroidopathy Associated with Hla-A29 Antigen and Immune Responsiveness to Retinal S-Antigen. Am J Ophthalmol. (1982) 94(2):147–158. 10.1016/0002-9394(82)90069-1

8. Herbort CP, Pavésio C, LeHoang P, Bodaghi B, Fardeau C, Kestelyn P, et al. Why birdshot retinochoroiditis should rather be called “HLA-A29 uveitis”? Br J Ophthalmol. (2017) 101(7):851–855. 10.1136/bjophthalmol-2016-309764

9. Kuiper JJW, Van Setten J, Ripke S, Van ‘T Slot R, Mulder F, Missotten T, et al. A genome-wide association study identifies a functional ERAP2 haplotype associated with birdshot chorioretinopathy. Hum Mol Genet. (2014) 23(22):6081–6087. 10.1093/hmg/ddu307

10. Kuiper JJW, Setten J van, Devall M, Cretu-Stancu M, Hiddingh S, Ophoff RA, et al. Functionally distinct ERAP1 and ERAP2 are a hallmark of HLA-A29-(Birdshot) Uveitis. Hum Mol Genet. (2018) 27(24):4333–4343. 10.1093/hmg/ddy319

11. Saveanu L, Carroll O, Lindo V, Del Val M, Lopez D, Lepelletier Y, et al. Concerted peptide trimming by human ERAP1 and ERAP2 aminopeptidase complexes in the endoplasmic reticulum. Nat Immunol. (2005) 6(7):689–697. 10.1038/ni1208

12. Andrés AM, Dennis MY, Kretzschmar WW, Cannons JL, Lee-Lin S-Q, Hurle B, et al. Balancing selection maintains a form of ERAP2 that undergoes nonsense-mediated decay and affects antigen presentation. PLoS Genet. (2010) 6(10):e1001157. 10.1371/journal.pgen.1001157

13. Mpakali A, Giastas P, Mathioudakis N, Mavridis IM, Saridakis E, Stratikos E. Structural basis for antigenic peptide recognition and processing by Endoplasmic reticulum (ER) aminopeptidase 2. J Biol Chem. (2015) 290(43):26021–26032. 10.1074/jbc.M115.685909

14. Giastas P, Mpakali A, Papakyriakou A, Lelis A, Kokkala P, Neu M, et al. Mechanism for antigenic peptide selection by endoplasmic reticulum aminopeptidase 1. Proc Natl Acad Sci. (2019) 116(52):26709–26716. 10.1073/pnas.1912070116

15. Evnouchidou I, Momburg F, Papakyriakou A, Chroni A, Leondiadis L, Chang SC, et al. The internal sequence of the peptide-substrate determines its N-Terminus trimming by ERAP1. PLoS One. (2008) 3(11). 10.1371/journal.pone.0003658

16. Birtley JR, Saridakis E, Stratikos E, Mavridis IM. The crystal structure of human endoplasmic reticulum aminopeptidase 2 reveals the atomic basis for distinct roles in antigen processing. Biochemistry. (2012) 51(1):286–295. 10.1021/bi201230p

17. Papakyriakou A, Reeves E, Beton M, Mikolajek H, Douglas L, Cooper G, et al. The partial dissociation of MHC class I-bound peptides exposes their N terminus to trimming by endoplasmic reticulum aminopeptidase 1. J Biol Chem. (2018) 293(20):7538–7548. 10.1074/jbc.RA117.000313

18. Chen H, Li L, Weimershaus M, Evnouchidou I, van Endert P, Bouvier M. ERAP1-ERAP2 dimers trim MHC I-bound precursor peptides; implications for understanding peptide editing. Sci Rep. (2016) 6:28902. 10.1038/srep28902

19. Mavridis G, Arya R, Domnick A, Zoidakis J, Makridakis M, Vlahou A, et al. A systematic re-examination of processing of MHCI-bound antigenic peptide precursors by endoplasmic reticulum aminopeptidase 1. J Biol Chem. (2020) 295(21):7193–7210. 10.1074/jbc.RA120.012976

20. Sanz-Bravo A, Martín-Esteban A, Kuiper JJW, García-Peydro M, Barnea E, Admon A, et al. Allele-specific alterations in the peptidome underlie the joint association of HLA-A*29:02 and endoplasmic reticulum aminopeptidase 2 (ERAP2) with birdshot chorioretinopathy. Mol Cell Proteomics. (2018) 17(8):1564–1577. 10.1074/mcp.RA118.000778

21. Alvarez-Navarro C, Martín-Esteban A, Barnea E, Admon A, López De Castro JA. Endoplasmic reticulum aminopeptidase 1 (ERAP1) polymorphism relevant to inflammatory disease shapes the peptidome of the birdshot chorioretinopathy-associated HLA-A*29:02 Antigen. Mol Cell Proteomics. (2015) 14(7):1770–1780. 10.1074/mcp.M115.048959

22. Georgiadis D, Mpakali A, Koumantou D, Stratikos E. Inhibitors of ER Aminopeptidase 1 and 2: From Design to Clinical Application. Curr Med Chem. (2019) 26(15):2715–2729. 10.2174/0929867325666180214111849

23. Anderson KS, Zeng W, Sasada T, Choi J, Riemer AB, Su M, et al. Impaired tumor antigen processing by immunoproteasome-expressing CD40-activated B cells and dendritic cells. Cancer Immunol Immunother. (2011) 60(6):857–867. 10.1007/s00262-011-0995-5

24. Hussong SA, Roehrich H, Kapphahn RJ, Maldonado M, Pardue MT, Ferrington DA. A novel role for the immunoproteasome in retinal function. Invest Ophthalmol Vis Sci. (2011) 52(2):714–723. 10.1167/iovs.10-6032

25. Hassan C, Kester MGD, Oudgenoeg G, de Ru AH, Janssen GMC, Drijfhout JW, et al. Accurate quantitation of MHC-bound peptides by application of isotopically labeled peptide MHC complexes. J Proteomics. (2014) 109:240–244. 10.1016/j.jprot.2014.07.009

26. Mulder A, Kardol MJ, Arn JS, Eijsink C, Franke MEI, Schreuder GMT, et al. Human monoclonal HLA antibodies reveal interspecies crossreactive swine MHC class I epitopes relevant for xenotransplantation. Mol Immunol. (2010) 47(4):809–815. 10.1016/j.molimm.2009.10.004

27. Brosch M, Yu L, Hubbard T, Choudhary J. Accurate and Sensitive Peptide Identification with Mascot Percolator. J Proteome Res. (2009) 8(6):3176–3181. 10.1021/pr800982s

28. Ritchie ME, Phipson B, Wu D, Hu Y, Law CW, Shi W, et al. limma powers differential expression analyses for RNA-sequencing and microarray studies. Nucleic Acids Res. (2015) 43(7):e47. 10.1093/nar/gkv007

29. Storey JD, Bass AJ, Dabney A, Robinson D. qvalue: Q-value estimation for false discovery rate control. R package version 2.14.1. (2019) https://github.com/idstorey/qvalue

30. Kammers K, Cole RN, Tiengwe C, Ruczinski I. Detecting Significant Changes in Protein Abundance. EuPA open proteomics. (2015) 7:11–19. 10.1016/j.euprot.2015.02.002

31. Sarkizova S, Klaeger S, Le PM, Li LW, Oliveira G, Keshishian H, et al. A large peptidome dataset improves HLA class I epitope prediction across most of the human population. Nat Biotechnol. (2020) 38(2):199–209. 10.1038/s41587-019-0322-9

32. Andreatta M, Lund O, Nielsen M. Simultaneous alignment and clustering of peptide data using a Gibbs sampling approach. Bioinformatics. (2013) 29(1):8–14. 10.1093/bioinformatics/bts621

33. McFerrin L. HDMD: Statistical Analysis Tools for High Dimension Molecular Data (HDMD). R package version 1.2. (2013) https://CRAN.R-project.org/package=HDMD

34. Abelin JG, Keskin DB, Sarkizova S, Hartigan CR, Zhang W, Sidney J, et al. Mass Spectrometry Profiling of HLA-Associated Peptidomes in Mono-allelic Cells Enables More Accurate Epitope Prediction. Immunity. (2017) 46(2):315–326. 10.1016/j.immuni.2017.02.007

35. Kim Y, Sidney J, Pinilla C, Sette A, Peters B. Derivation of an amino acid similarity matrix for peptide: MHC binding and its application as a Bayesian prior. BMC Bioinformatics. (2009) 10:394. 10.1186/1471-2105-10-394

36. Goslee SC, Urban DL. The ecodist Package for Dissimilarity-based Analysis of Ecological Data. J. Stat. Software. (2007) 1(7). 10.18637/jss.v022.i07

37. Hahsler M, Piekenbrock M, Doran D. dbscan: Fast Density-Based Clustering with R. J Stat Software. (2019) 1(1). 10.18637/jss.v091.i01

38. Hennig C. fpc: Flexible Procedures for Clustering. R package version 2.2-5. (2020) https://CRAN.R-project.org/package=fpc

39. Wagih O. ggseqlogo: A ‘ggplot2’ Extension for Drawing Publication-Ready Sequence Logos. R package version 0.1. (2017) https://CRAN.R-project.org/package=ggseqlogo

40. Osorio D, Rondón-Villarreal P, Torres Sáez R. Peptides: A Package for Data Mining of Antimicrobial Peptides. R J. (2015) 7:4–14. 10.32614/RJ-2015-001

41. Ogle DH, Wheeler P, Dinno A. FSA: Fisheries Stock Analysis. R package version 0.8.30. (2020) https://github.com/droglenc/FSA.

42. Gfeller D, Guillaume P, Michaux J, Pak H-S, Daniel RT, Racle J, et al. The Length Distribution and Multiple Specificity of Naturally Presented HLA-I Ligands. J Immunol. (2018) 201(12):370516. 10.4049/jimmunol.1800914

43. René C, Lozano C, Villalba M, Eliaou J-F. 5’ and 3’ untranslated regions contribute to the differential expression of specific HLA-A alleles. Eur J Immunol. (2015) 45(12):3454–3463. 10.1002/eji.201545927

44. López de Castro JA, Alvarez-Navarro C, Brito A, Guasp P, Martín-Esteban A, Sanz-Bravo A. Molecular and pathogenic effects of endoplasmic reticulum aminopeptidases ERAP1 and ERAP2 in MHC-I-associated inflammatory disorders: Towards a unifying view. Mol Immunol. (2016) 77:193–204. 10.1016/j.molimm.2016.08.005

45. Rao X, Hoof I, Costa AICAF, van Baarle D, Kesmir C. HLA class I allele promiscuity revisited. Immunogenetics. (2011) 63(11):691–701. 10.1007/s00251-011-0552-6

46. Chowell D, Krishna S, Becker PD, Cocita C, Shu J, Tan X, et al. TCR contact residue hydrophobicity is a hallmark of immunogenic CD8+ T cell epitopes. Proc Natl Acad Sci U S A. (2015) 112(14):E1754–1762. 10.1073/pnas.1500973112

47. Riley TP, Keller GLJ, Smith AR, Davancaze LM, Arbuiso AG, Devlin JR, et al. Structure Based Prediction of Neoantigen Immunogenicity. Front Immunol. (2019) 10:2047. 10.3389/fimmu.2019.02047

48. Lim YW, Chen-Harris H, Mayba O, Lianoglou S, Wuster A, Bhangale T, et al. Germline genetic polymorphisms influence tumor gene expression and immune cell infiltration. Proc Natl Acad Sci USA. (2018) 115(50):E11701–11710. 10.1073/pnas.1804506115

49. Coles CH, McMurran C, Lloyd A, Hock M, Hibbert L, Raman MCC, et al. T Cell Receptor interactions with Human Leukocyte Antigen govern indirect peptide selectivity for the cancer testis antigen MAGE-A4. J Biol Chem. (2020) 295:11486–11494. 10.1074/jbc.RA120.014016

50. Márquez A, Cordero-Coma M, Martín-Villa JM, Gorroño-Echebarría MB, Blanco R, Díaz Valle D, et al. New insights into the genetic component of non-infectious uveitis through an Immunochip strategy. J Med Genet. (2017) 54(1):38–46. 10.1136/jmedgenet-2016-104144

51. Apps R, Meng Z, Del Prete GQ, Lifson JD, Zhou M, Carrington M. Relative Expression Levels of the HLA Class-I Proteins in Normal and HIV-Infected Cells. J Immunol. (2015) 1403234. 10.4049/jimmunol.1403234

52. Kuiper JJW, Venema WJ. HLA-A29 and Birdshot Uveitis: Further Down the Rabbit Hole. Front Immunol. (2020) 11:599558. 10.3389/fimmu.2020.599558

53. Elahi S, Herbort CPJ. Vogt-Koyanagi-Harada Disease and Birdshot Retinochoroidopathy, Similarities and Differences: A Glimpse into the Clinicopathology of Stromal Choroiditis, a Perspective and a Review. Klin Monbl Augenheilkd. (2019) 236(4):492–510. 10.1055/a-0829-6763

54. Papadia M, Herbort CP. New concepts in the appraisal and management of birdshot retinochoroiditis, a global perspective. Int Ophthalmol. (2015) 35(2):287–301. 10.1007/s10792-015-0046-x

55. Hassman L, Warren M, Huxlin KR, Chung MM, Xu L. Evidence of melanoma immunoreactivity in patients with Birdshot retinochoroidopathy. Invest Ophthalmol Vis Sci. (2017) 58(8):5745.

56. Chen L, Shi H, Koftori D, Sekine T, Nicastri A, Ternette N, et al. Identification of an Unconventional Subpeptidome Bound to the Behçet’s Disease-associated HLA-B*51:01 that is Regulated by Endoplasmic Reticulum Aminopeptidase 1 (ERAP1). Mol Cell Proteomics. (2020) 19(5):871–883. 10.1074/mcp.ra119.001617

57. Guasp P, Lorente E, Martín-Esteban A, Barnea E, Romania P, Fruci D, et al. Redundancy and Complementarity between ERAP1 and ERAP2 revealed by their effects on the behcet’s disease-associated HLA-B*51 peptidome. Mol Cell Proteomics. (2019) 18(8):1491–1510. 10.1074/mcp.RA119.001515

58. Heterozygosity of the 721.221-B*51:01 Cell Line Used in the Study by Guasp et (Arthritis Rheumatol, February 2016). Arthritis Rheumatol. (2017) 69(3):686. 10.1002/art.40073

## References Supplemental Info

Alvarez-Navarro C, Martín-Esteban A, Barnea E, Admon A, López de Castro JA. Endoplasmic Reticulum Aminopeptidase 1 (ERAP1) Polymorphism Relevant to Inflammatory Disease Shapes the Peptidome of the Birdshot Chorioretinopathy-Associated HLA-A*29:02 Antigen. Mol Cell Proteomics. 2015 Jul;14(7): 1770–80.

Brosch M, Yu L, Hubbard T, Choudhary J. Accurate and sensitive peptide identification with Mascot Percolator. J Proteome Res. 2009 Jun;8(6):3176–81.

Kammers K, Cole RN, Tiengwe C, Ruczinski I. Detecting Significant Changes in Protein Abundance. EuPA Open Proteom. 2015 Jun;7:11–19.

Sanz-Bravo A, Martín-Esteban A, Kuiper JJW, García-Peydró M, Barnea E, Admon A, López de Castro JA. Allele-specific Alterations in the Peptidome Underlie the Joint Association of HLA-A*29:02 and Endoplasmic Reticulum Aminopeptidase 2 (ERAP2) with Birdshot Chorioretinopathy. Mol Cell Proteomics. 2018 Aug;17(8):1564–1577.

Sarkizova S, Klaeger S, Le PM, Li LW, Oliveira G, Keshishian H, Hartigan CR, Zhang W, Braun DA, Ligon KL, Bachireddy P, Zervantonakis IK, Rosenbluth JM, Ouspenskaia T, Law T, Justesen S, Stevens J, Lane WJ, Eisenhaure T, Lan Zhang G, Clauser KR, Hacohen N, Carr SA, Wu CJ, Keskin DB. A large peptidome dataset improves HLA class I epitope prediction across most of the human population. Nat Biotechnol. 2020 Feb;38(2):199–209.

